# EPIC: inferring relevant cell types for complex traits by integrating genome-wide association studies and single-cell RNA sequencing

**DOI:** 10.1101/2021.06.09.447805

**Authors:** Rujin Wang, Dan-Yu Lin, Yuchao Jiang

## Abstract

More than a decade of genome-wide association studies (GWASs) have identified genetic risk variants that are significantly associated with complex traits. Emerging evidence suggests that the function of trait-associated variants likely acts in a tissue- or cell-type-specific fashion. Yet, it remains challenging to prioritize trait-relevant tissues or cell types to elucidate disease etiology. Here, we present EPIC (cEll tyPe enrIChment), a statistical framework that relates large-scale GWAS summary statistics to cell-type-specific gene expression measurements from single-cell RNA sequencing (scRNA-seq). We derive powerful gene-level test statistics for common and rare variants, separately and jointly, and adopt generalized least squares to prioritize trait-relevant cell types while accounting for the correlation structures both within and between genes. Using enrichment of loci associated with four lipid traits in the liver and enrichment of loci associated with three neurological disorders in the brain as ground truths, we show that EPIC outperforms existing methods. We apply our framework to multiple scRNA-seq datasets from different platforms and identify cell types underlying type 2 diabetes and schizophrenia. The enrichment is replicated using independent GWAS and scRNA-seq datasets and further validated using PubMed search and existing bulk case-control testing results.

## Background

Many years of genome-wide association studies (GWASs) have yielded genetic risk variants associated with complex traits and human diseases. Emerging evidence suggests that the function of trait-associated variants likely acts in a tissue- or cell-type-specific fashion ^1, 2, 3, 4, 5^. Recent advances in single-cell RNA sequencing (scRNA-seq) ^6, 7, 8^ enable characterization of cell-type-specific gene expression and provide an unprecedented opportunity to systematically investigate the cell-type-specific enrichment of GWAS polygenic signals ^9, 10, 11, 12^. There is a pressing need to develop a statistically rigorous and computationally scalable analytical framework to integrate large-scale genome-wide association studies (e.g., the UK Biobank ^13^) and high-dimensional scRNA-seq efforts (e.g., the Human Cell Atlas ^14^). Such an integrative analysis helps elucidate the underlying cell-type-specific disease etiology and prioritize important functional variants.

Several methods ^15, 16, 17, 18, 19^ have been developed to integrate scRNA-seq data with GWAS summary statistics to prioritize trait-relevant cell types. One set of methods, including RolyPoly ^15^ and LDSC-SEG ^17^, develops models on the single-nucleotide polymorphism (SNP) level and derives SNP-wise annotations from the transcriptomic data. RolyPoly adopts a polygenic model, and the effect sizes of all SNPs associated with a gene have a covariance that is a linear combination of the gene expressions across all cell types. RolyPoly, therefore, captures the effect of the cell-type-specific gene expression on the covariance of GWAS effect sizes. LDSC-SEG also constructs SNP annotations from cell-type-specific gene expressions and then carries out a one-sided test using the stratified LD score regression framework ^17, 20, 21, 22^. It tests whether trait heritability is enriched in regions surrounding genes that have the highest specific expression in a given cell type.

Another set of methods, such as CoCoNet ^18^ and MAGMA ^9, 16, 23, 24^, does not devise the SNP-level framework. These methods first derive gene-level association statistics since this more naturally copes with the gene-level expression measurements; they then prioritize risk genes in a specific cell type. Specifically, CoCoNet models gene-level association statistics as a function of the cell-type-specific adjacency matrices inferred from gene expression studies. While CoCoNet is the first method to evaluate the gene co-expression networks, its rank-based method does not allow hypothesis testing due to the strong correlation among gene co-expression patterns constructed from different cell types. Like CoCoNet, MAGMA ^16^ and MAGMA-based approaches ^9, 23, 24^ also begin by combining SNP-level GWAS summary statistics into gene-level statistics. This step is followed by a second “gene-property” analysis, where the cell-type-specific gene expressions are regressed against the genes’ GWAS test statistics. The various versions of the methods adopt different ways to select genes, transform the outcome and predictor variables, and include different sets of additional covariates ^9, 23, 24^. While MAGMA-based methods have been successfully used in several studies ^25, 26, 27^, Yurko et al. ^28^ examined the statistical foundation of MAGMA, and they identified an issue: type I error rate is inflated because the method incorrectly uses the Brown’s approximation when combining the SNP-level *p* -values. In addition to this problem, we noticed that the MAGMA’s implementation uses squared correlations between SNPs, which masks the true LD structure.

When modeling on the gene level, one needs to account for the gene-gene correlations. RolyPoly ignores proximal gene correlations but implements a block bootstrapping procedure as a correction. MAGMA approximates the gene-gene correlations as the correlations between the model sum of squares from the second-step gene-property analysis. However, the gene-gene correlation of the effect sizes should be a function of the LD scores (i.e., the correlations between the SNPs within the genes). CoCoNet does not take account of this either, instead using LD information only to calculate the gene-level effect sizes and assuming that gene-gene covariance is a function solely of gene co-expression. A statistically rigorous and computationally efficient method to derive the gene-gene correlation structure while incorporating the SNP-level LD information is needed.

These existing methods either focus on common variants (e.g., RolyPoly and LDSC-SEG) or do not differentiate between common and rare variants (e.g., MAGMA with only summary statistics) due to the limited statistical power for rare variants. While methods for rare-variant association analysis have been developed (e.g., sequence kernel association test ^29^ and burden test ^30^), to our best knowledge, no methods are currently available to detect cell-type-specific enrichment of GWAS risk loci using summary statistics for both common and rare variants.

Here, we propose EPIC, a statistical framework to identify trait-relevant cell types by integrating GWAS summary statistics and cell-type-specific gene expression profiles from scRNA-seq. We adopt gene-based generalized least squares to identify enrichment of risk loci. For the prioritized cell types, EPIC further carries out a gene-specific influence analysis to identify significant genes. We demonstrate EPIC on multiple tissue-specific bulk RNA-seq and scRNA-seq datasets, along with GWAS summary statistics of four lipid traits, three neuropsychiatric disorders, and type 2 diabetes, and successfully replicate and validate the prioritized tissues and cell types. Together, EPIC’s integrative analysis of cell-type-specific expressions and GWAS polygenic signals help to elucidate the underlying cell-type-specific disease etiology and prioritize important functional variants. EPIC is compiled as an open-source R package available at https://github.com/rujinwang/EPIC.

## Results

### Overview of methods

The goal of EPIC is to identify disease- or trait-relevant cell types. An overview of the framework is outlined in Figure 1. EPIC takes as input single-variant summary statistics from GWAS, which is used to aggregate SNP-level associations into genes, and gene expression datasets from scRNA-seq data. An external reference panel is adopted to account for the linkage disequilibrium (LD) between SNPs and genes. We first perform gene-level testing based on GWAS summary statistics from the single-variant analysis. The multivariate statistics for both common and rare variants can be recovered using covariance of the single-variant test statistics, which can be estimated from either the participating study or from a public database. We then develop a gene-based regression framework that can prioritize trait-relevant cell types from gene-level test statistics and cell-type-specific omics profiles while accounting for gene-gene correlations due to LD. The underlying hypothesis is that if a particular cell type influences a trait, then more of the GWAS polygenic signals would be concentrated in genes with greater cell-type-specific gene expression. For significantly enriched cell type(s), we further carry out a gene-specific influence analysis to identify genes that are highly influential in leading to the significance of the prioritized cell type. Refer to the Methods section for methodological and algorithmic details.

**Figure 1.**
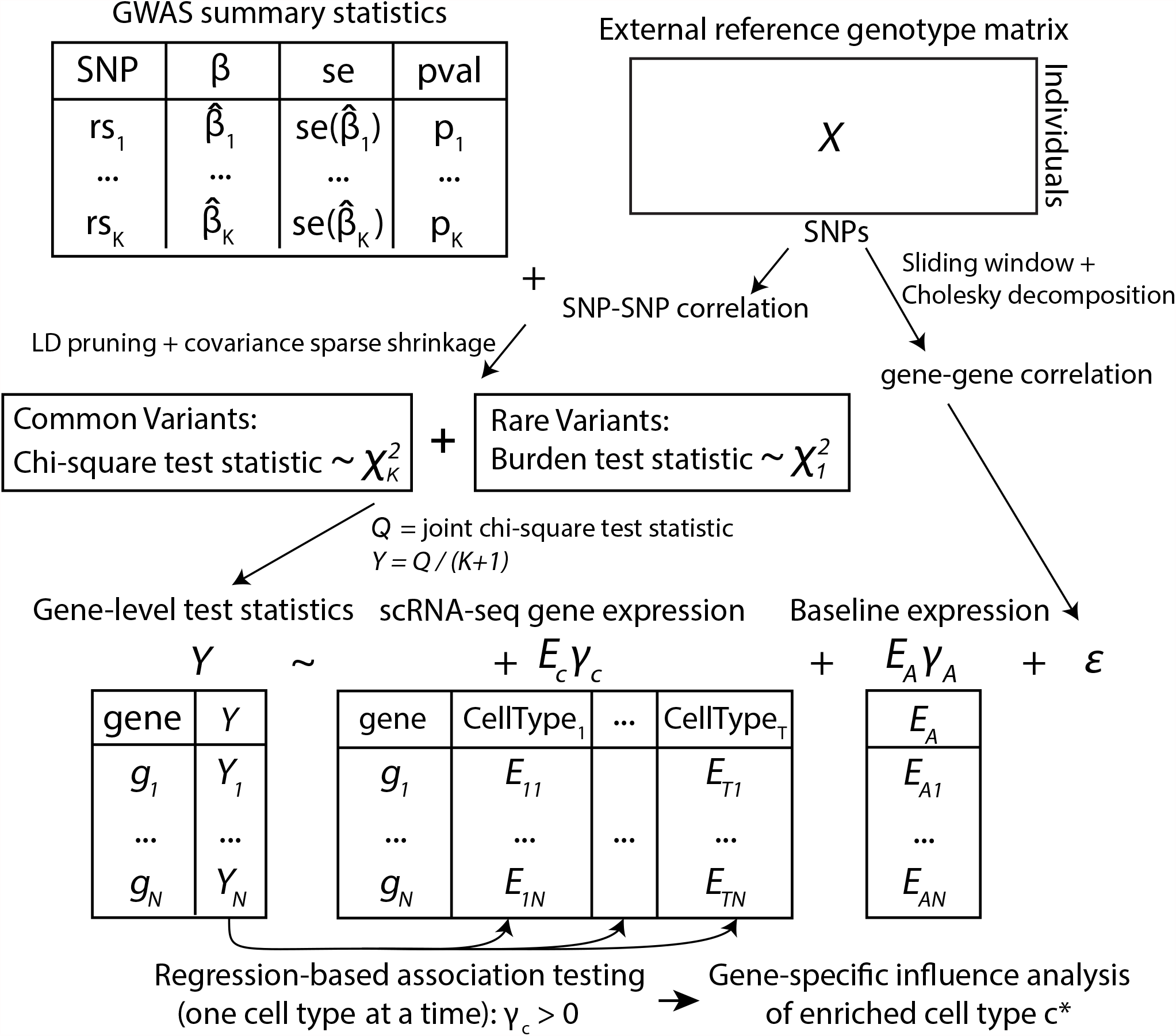
Overview of EPIC framework. EPIC starts from GWAS summary statistics and an external reference panel to account for LD structure. To ensure that the correlation matrix is well-conditioned, EPIC adopts the POET estimators to obtain a sparse shrinkage correlation matrix. EPIC performs LD pruning, computes the gene-level chi-square statistics for common variants, and calculates burden test statistics for rare variants. EPIC then integrates gene-level association statistics with transcriptomic profiles and prioritizes trait-relevant cell types using a regression-based framework while accounting for the gene-gene correlation structure.

### Inferring trait-relevant tissues using tissue-specific RNA-seq

As a proof of concept, we started our analysis with GWAS summary statistics for eight diseases and traits (four lipid traits ^31^, three neuropsychiatric disorders ^32, 33, 34^, and T2Db ^35^) and tissue-specific transcriptomic profiles from the Genotype-Tissue Expression project (GTEx) v8 ^36^ (Supplementary Table S1). The GTEx consortium consists of bulk-tissue gene expression measurements of 17,382 samples from 54 tissues across 980 postmortem donors; after sample-specific quality controls, we obtained gene expression profiles of 45 tissues, averaged across samples (Supplementary Table S1). For subsequent analyses, we focused on a set of 8,708 genes with tissue specificity scores greater than 5 (Supplementary Note S2).

We first performed the gene-level chi-square association test with the shrinkage estimators and sliding-window approach. In Supplementary Table S2, we summarized a list of genes that have been previously reported to be associated with traits ^31, 32, 33, 34, 35^; for these sets of well characterized trait-associated genes, EPIC returned much more significant *p* -values compared to MAGMA. On the genome-wide scale, the quantile-quantile plots of gene-level *p* -values demonstrated EPIC’s elevated power (Supplementary Figure S1), and EPIC detected more significant genes than MAGMA after Bonferroni correction (Supplementary Figure S2). Importantly, we further reported gene-level association testing results for a set of housekeeping genes ^37^ and demonstrated that, while powerful, EPIC also controlled for type I error (Supplementary Figure S3).

We next applied EPIC to identify the trait-relevant tissues by performing tissue-specific regression for each trait, with results shown in Figure 2, Figure 3A, and Figure 4A. All four lipid traits are significantly enriched in the liver (Figure 2), which plays a key role in lipid metabolism. The small intestine was marginally enriched for TC – it has been shown that the small intestine plays an important role in cholesterol regulation and metabolism ^38, 39^. In addition, the adipose tissues, which have also been shown to regulate lipid metabolism ^40, 41^, were identified as being significantly enriched by both EPIC and MAGMA. Both LDSC-SEG and RolyPoly suffer from low power, although the liver was one of the top-ranked tissues for the lipid traits.

**Figure 2.**
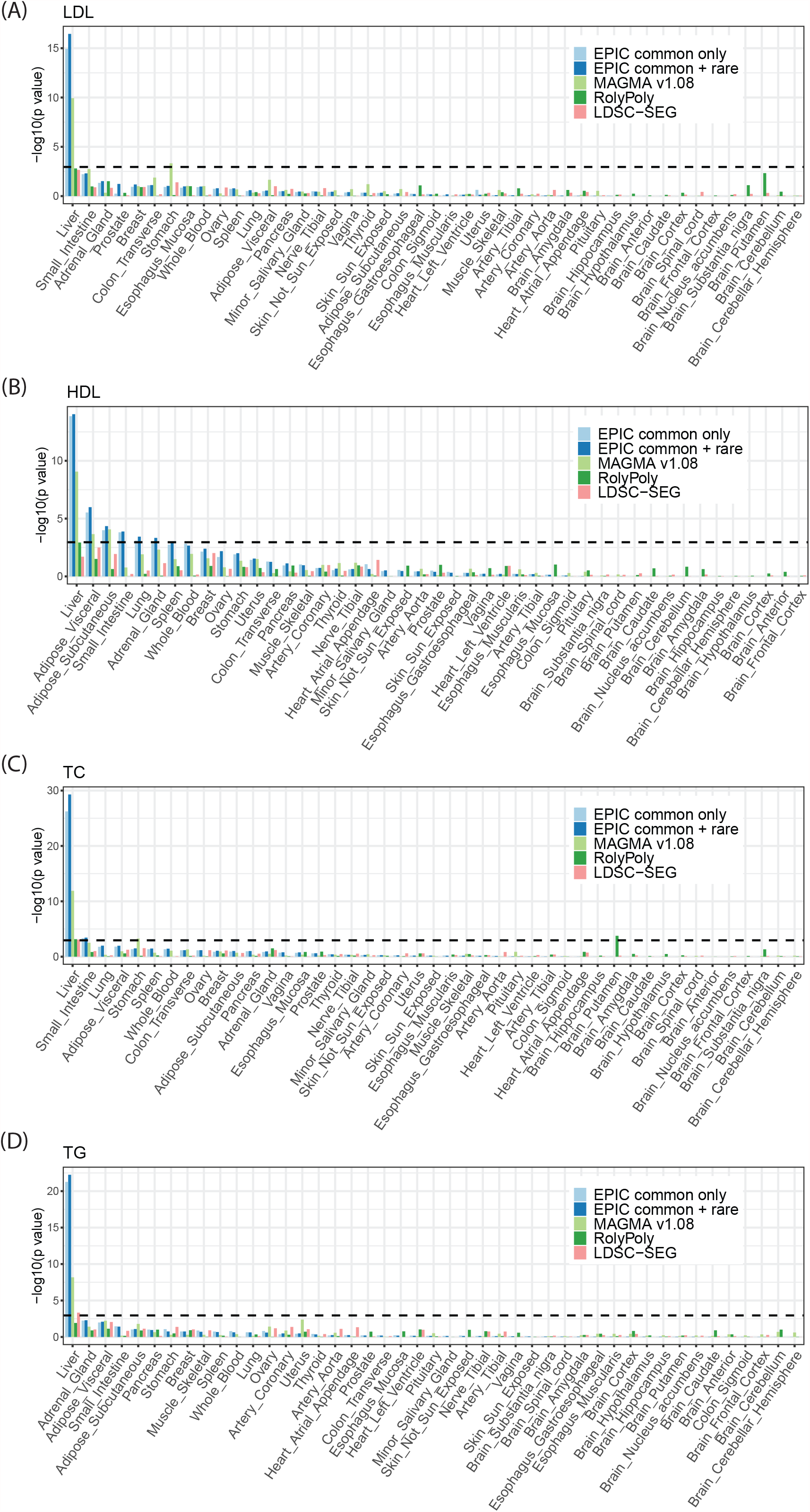
Tissue enrichment for four lipid traits using GTEx bulk RNA-seq data. (A) LDL; (B) HDL; (C) TC; and (D) TG. The dashed line is the Bonferroni-corrected *p*-value threshold. EPIC achieved higher power while controlling for false positives compared to other existing methods.

**Figure 3.**
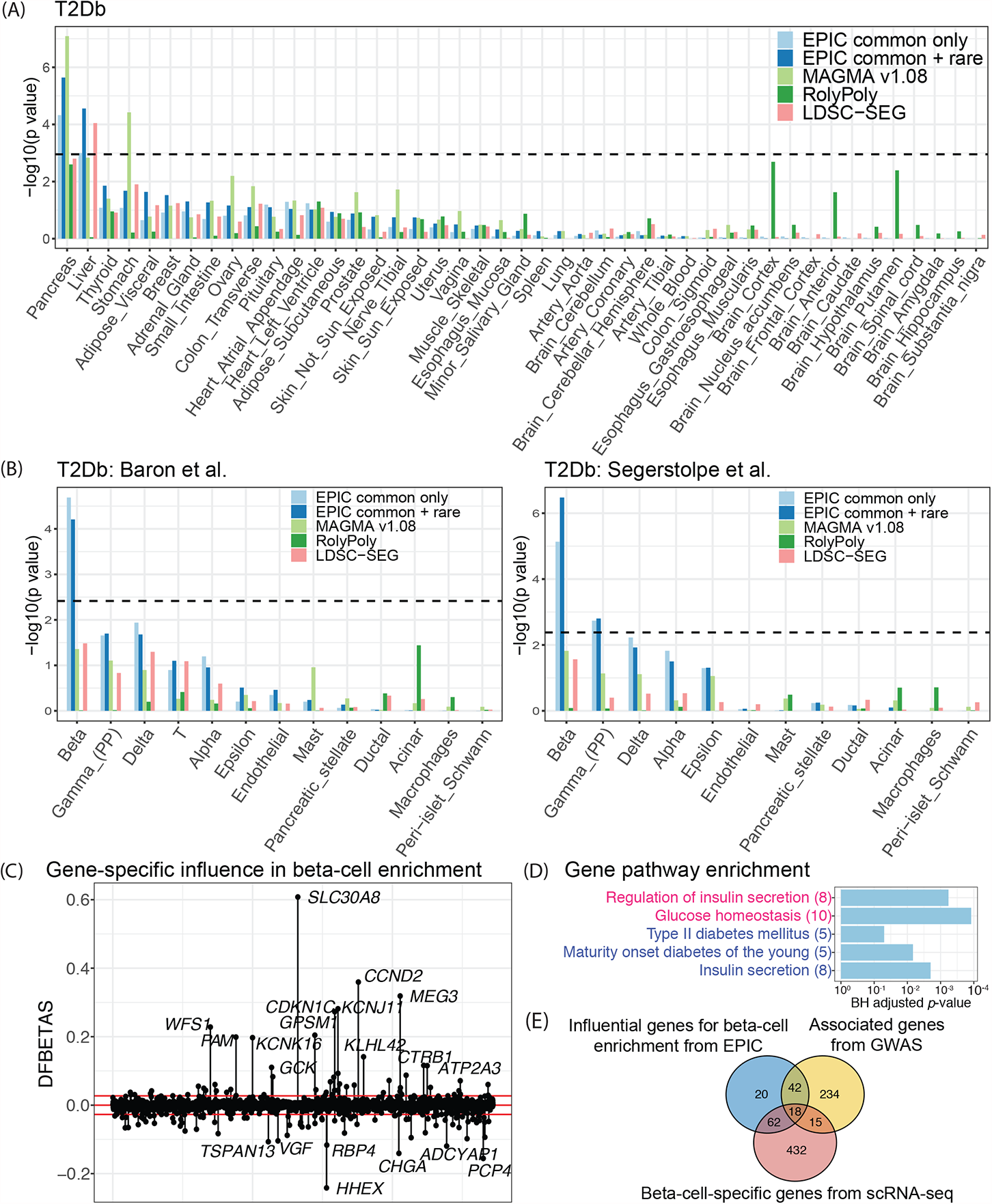
Cell-type-specific enrichment of T2Db risk loci. (A) T2Db-relevant tissue identification using GTEx tissue-specific RNA-seq data. (B) T2Db-relevant cell type identification using scRNA-seq data of human pancreatic islets. The dashed line is the Bonferroni-corrected *p*-value threshold. (C) Gene-specific influence analysis for the significantly enriched beta cells. DFBETA measures the difference in the estimated coefficients in the gene-property analysis with and without each gene. Red lines are the size-adjusted cutoffs 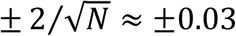, where *N* is the number of genes. (D) Gene pathway enrichment analysis using highly influential genes. KEGG pathways (pink) and GO biological processes (blue) related to T2Db are significantly enriched. (E). Venn diagram of the significant genes from the beta-cell-specific influential analysis by EPIC, gene-level association testing from GWAS, and nonparametric testing of cell-type-specific expression from scRNA-seq. The highly influential genes that lead to the enrichment of beta cells are significantly associated with the trait and/or specifically expressed in the cell type.

**Figure 4.**
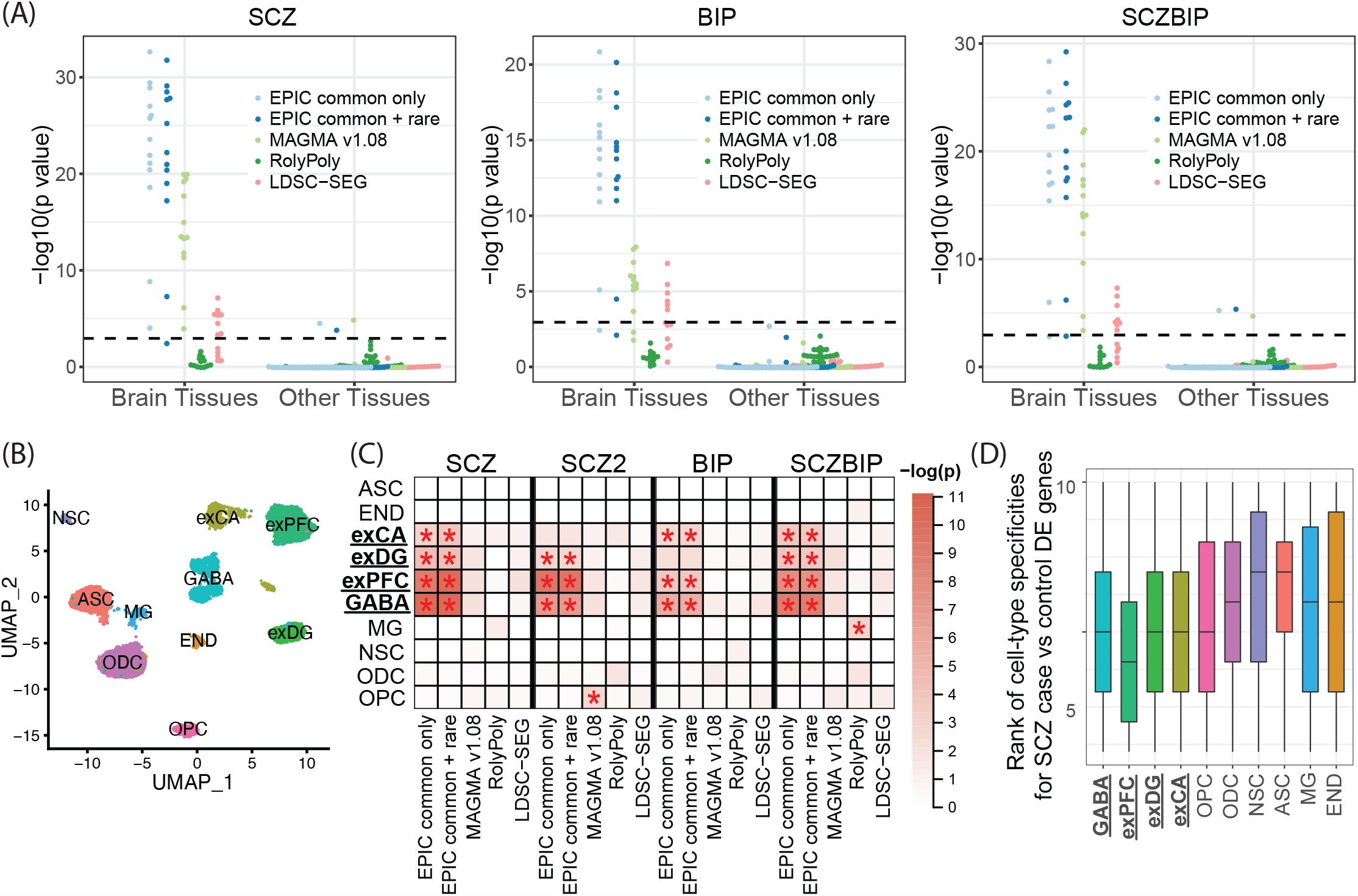
Cell-type-specific enrichment for three neuropsychiatric disorders. (A) Beeswarm plot of –log10(*p*-value) from the tissue enrichment analysis using GTEx bulk RNA-seq data. The dashed line is Bonferroni corrected *p*-value threshold (0.05/45). (B) UMAP embedding of 14,137 single cells from five donors. (C) Heatmap of –log10(*p*-value) from the cell-type enrichment analysis using GTEx scRNA-seq brain data. Bonferroni-significant results are marked with red asterisks (*p*<0.05/10). GABA: GABAergic interneurons; exPFC: excitatory glutamatergic neurons in the prefrontal cortex; exDG: excitatory granule neurons from the hippocampal dentate gyrus region; exCA: excitatory pyramidal neurons in the hippocampal Cornu Ammonis region; OPC: oligodendrocyte precursor cells; ODC: oligodendrocytes; NSC: neuronal stem cells; ASC: astrocytes; MG: microglia cells; END: endothelial cells. (D) Boxplots of gene specificity ranks across ten brain cell types for differentially expressed genes from SCZ case-control studies.

Neuropsychiatric disorders exhibited strong brain-specific enrichments, as expected. The frontal cortex of the brain was detected as being the most strongly enriched for SCZ, BIP, and SCZBIP (Figure 4A). The pituitary also demonstrated strong enrichment signals with SCZ and SCZBIP, while the spinal cord was found to be an irrelevant tissue with these three neuropsychiatric disorders. In comparison, LDSC-SEG identified part of the brain tissues as trait-relevant, while RolyPoly failed to return enrichment in any of the brain tissues (Figure 4A).

As a final proof of concept, we sought to infer T2Db-relevant tissue(s) using the tissue-specific gene expression data GTEx. The pancreas and the liver were prioritized as the T2Db-relevant tissues by EPIC, while MAGMA yielded significant results in the pancreas as well as the stomach (Figure 3A). RolyPoly identified the pancreas as the second most relevant tissue; LDSC-SEG reported the liver as the only significantly enriched tissue (Figure 3A).

Notably, we have thus far focused on carrying out the enrichment analysis using common variants only or using common and rare variants combined. For rare variants alone, EPIC successfully identified liver and brain as the top-rank tissue for the lipid traits and the neuropsychiatric disorders, respectively (Supplementary Table S3), although it is generally underpowered. This is possibly due to GWAS being underpowered to detect rare-variant associations (Supplementary Table S2) and the expression profiles of rare variants being hard to be recapitulated by a single-cell reference. Nevertheless, while current studies only reported common variants that were consistently mapped to a subset of brain cell types for neuropsychiatric disorders ^9, 23^, EPIC offers a statistical framework to identify cell-type-specific enrichment signals attributed to both common and rare variants, separately and jointly.

For validation, we adopted a similar strategy as proposed by Shang et al. ^18^ – we carried out a PubMed search, resorting to previous literature studying the trait of interest in relation to a particular tissue or cell type. Specifically, we counted the number of previous publications using the keyword pairs of trait and tissue/cell type and calculated the Spearman’s rank correlations between the number of publications and EPIC’s tissue-/cell-type-specific *p* -values (Figure 5). Across all traits, we found strong positive correlations between EPIC’s enrichment results and PubMed search results (Figure 5A).

**Figure 5.**
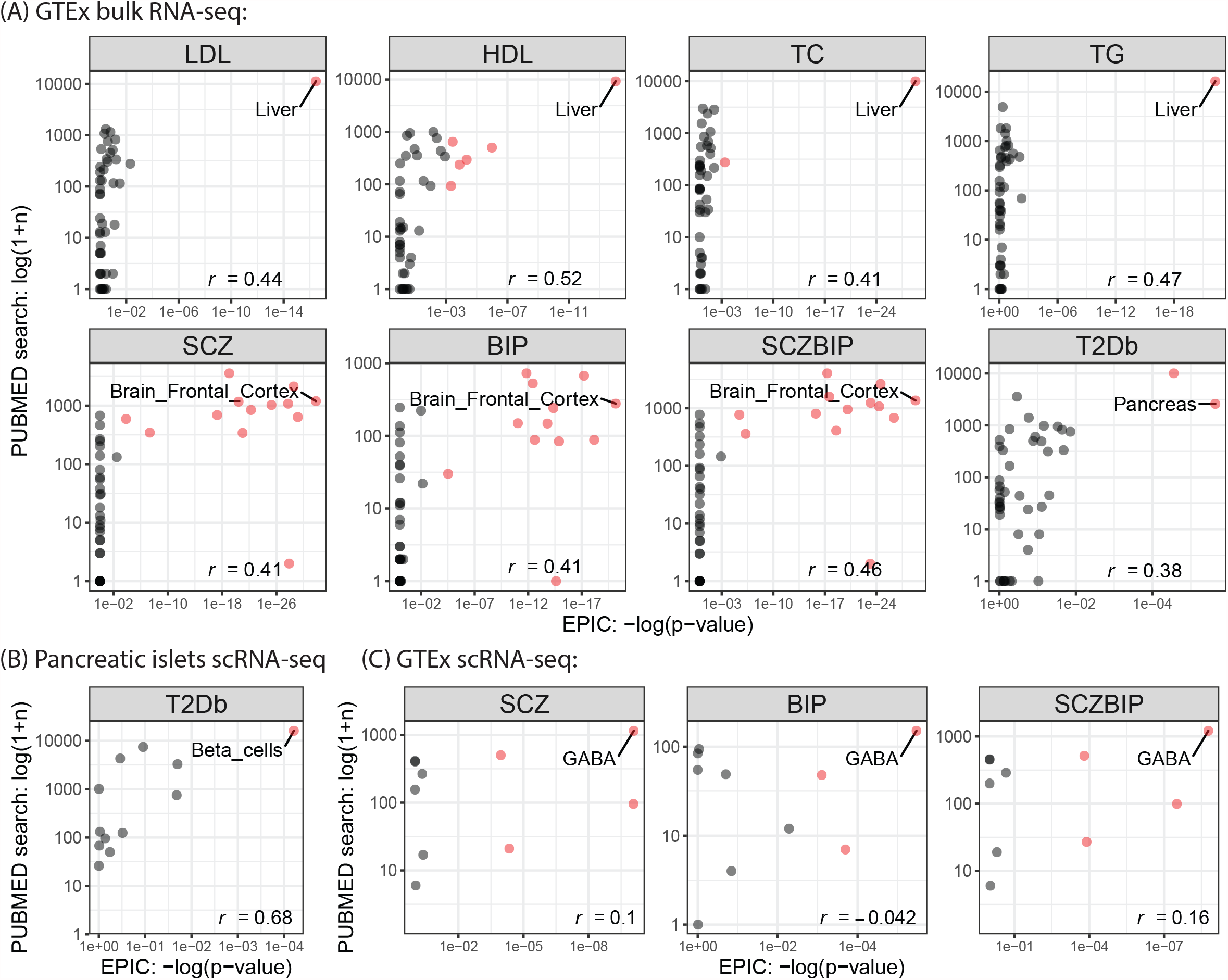
Correlations of tissue/cell type ranks from enrichment analysis and PubMed Search. Spearman correlations are calculated between the PubMed search and EPIC’s results. Trait-relevant tissues/cell types with statistical significance after Bonferroni correction are highlighted in red, where the top-ranking tissues/cell types are labeled.

### Inferring relevant cell types for T2Db by scRNA-seq data of pancreatic islets

We next analyzed pancreatic islet scRNA-seq data to identify trait-relevant cell types for T2Db. To assess reproducibility, EPIC was separately applied to two scRNA-seq datasets consisting of multiple endocrine cell types (Supplementary Table S1 and Supplementary Figure S4). The scRNA-seq data were generated using two different protocols: the SMART-Seq2 protocol on six healthy donors from Segerstolpe et al. ^8^ and the InDrop protocol on three healthy individuals from Baron et al. ^6^. In both datasets, beta cells were identified as the trait-relevant cell types by EPIC (Figure 3B). Enrichment of beta cells is used as a gold standard for benchmark, in that beta cells produce and release insulin but are dysfunctional and gradually lost in T2Db ^42, 43^. We also found that gamma cells were marginally associated with T2Db in the Segerstolpe dataset – pancreatic polypeptide, which is produced by gamma cells, is known to play a critical role in endocrine pancreatic secretion regulation ^44, 45, 46^. As a comparison, neither MAGMA nor LDSC-SEG detected significant enrichment in beta cells, even though the enrichment was top-ranked. RolyPoly, on the other hand, did not report any enrichment of the beta cells compared to the other types of cells.

To additionally validate the beta-cell enrichment, we carried out the PubMed search using the trait-and-cell-type pairs as keywords and showed that the cell-type ranks obtained from EPIC’s beta-cell-specific *p*-values were highly consistent with those from the PubMed search results (Figure 5B). Together, we demonstrate the effectiveness of EPIC in identifying trait-relevant cell types using scRNA-seq datasets generated by different protocols.

To identify specific genes that drive the significant enrichment in beta cells, we carried out the gene-specific influence test as outlined in Methods and identified 142 highly influential genes (Figure 3C). We performed KEGG pathway analysis and Gene Ontology (GO) biological process enrichment analysis using the DAVID bioinformatics resources ^47, 48^. Beta-cell-specific influential genes are enriched in GO terms including glucose homeostasis and regulation of insulin secretion, as well as KEGG pathways including insulin secretion, maturity onset diabetes, etc. (Figure 3D). Compared to the set of genes that are significantly associated with T2Db from GWAS and the set of genes that are specifically expressed in beta cells from scRNA-seq, the set of highly influential genes that led to the enrichment of beta cells are, to a great degree, highly associated with the trait and/or specifically expressed in the cell type of interest (Figure 3E). However, the Venn diagram suggests that they do not perfectly overlap, especially across all three sets. The influential analysis by EPIC helps prioritize trait-relevant genes in a cell-type-specific manner.

### Inferring relevant cell types for neuropsychiatric disorders by scRNA-seq data of human brain

To further test EPIC in a more complex tissue, we sought to prioritize trait-relevant cell types in the brain. While the brain tissues are significantly enriched using the GTEx bulk-tissue RNA-seq data (Figure 4A), the relevant cell types in the brain for neuropsychiatric disorders are not as well defined and studied. We obtained droplet-based scRNA-seq data ^7^, generated on frozen adult human postmortem tissues from the GTEx project (Supplementary Table S1), to infer the relevant cell types. After pre-processing and stringent quality controls, the scRNA-seq data contains gene expression profiles of 17,698 genes across 14,137 single cells collected from the human hippocampus and prefrontal cortex tissues. The cells belong to ten cell types (Figure 4B), and we focused on the top 8,000 highly variable genes for subsequent analyses.

We evaluated EPIC’s cell-type-specific enrichment results and found that all three neuropsychiatric disorders were significantly enriched in GABAergic interneurons (GABA), excitatory glutamatergic neurons from the prefrontal cortex (exPFC), and excitatory pyramidal neurons in the hippocampal CA region (exCA). Excitatory granule neurons from the hippocampal dentate gyrus region (exDG) were identified as relevant cell types for SCZ and SCZBIP (Figure 4C). EPIC successfully replicated the previously reported association of neuropsychiatric disorders with interneurons and excitatory pyramidal neurons ^9, 23^.

We employed three strategies to validate the trait-relevant cell types for the neuropsychiatric disorders. First, we again found positive Spearman correlations with PubMed search results and EPIC’s enrichment results for SCZ and SCZBIP (Figure 5C). Second, we adopted additional independent GWAS summary statistics for SCZ (SCZ2) ^49^ (Supplementary Table S1) and observed highly concordant enrichment results between SCZ and SCZ2 (Figure 4C). Third, we tested whether genes that are upregulated/downregulated for SCZ were enriched in the identified cell types to additionally implicate cell types involved in SCZ. Specifically, we performed differential expression (DE) analysis from an independent case-control study of SCZ using bulk RNA-seq ^50^, retaining 287 significant DE genes that also overlap the scRNA-seq data (Supplementary Figure S5). We reasoned that, if SCZ-relevant risk loci were enriched in a particular cell type, genes that are differentially expressed between SCZ cases and controls would demonstrate greater cell-type specificity in this cell type. We calculated cell-type specificities using the set of DE genes and observed GABA, exCA, exDG, and exPFC were the top four cell types with the lowest gene-specificity ranks (Figure 4D). Using three different strategies by querying external databases and adopting additional and orthogonal datasets, we validated the trait-cell-type relevance results.

## Methods

### Gene-level associations for common variants

Let *β* = (*β*_1_, ⋯, *β*_*K*_)^*T*^ be the effect sizes of *K* common variants within a gene of interest. Let 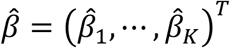 be the estimator for *β*, with corresponding standard error 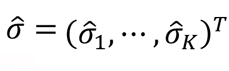. Let 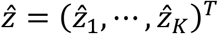 be the z -scores, where 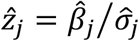 is the standard-normal statistic for testing the null hypothesis of no association for SNP *j*. We approximate the correlation matrix of 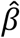 (equivalent to the covariance matrix of 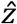) by the LD matrix *R* = {*R*_*jl*_; *j, l* = 1, *…, K*}, where *R*_*jl*_ is the Pearson correlation between SNP *j* and SNP *l*. We further define 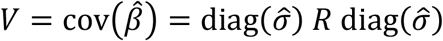 as the covariance matrix of 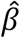. Under the null hypothesis of *β* = 0, the estimator 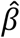 is *K*-variate normal with mean 0 and covariance matrix *V*. To perform gene-level association testing for common variants, we construct a simple and powerful chi-square statistic for testing the null hypothesis of *β* = 0:

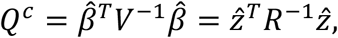

which has the 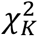 distribution under the null. The correlation matrix *R* can be estimated from either the participating study or a publicly available reference panel. In this study, we utilize the 1000 Genomes Project European panel ^51^, which comprises genotypes of ∼500 European individuals across ∼23 million SNPs.

An effective chi-square test described above requires the covariance matrix to be well-conditioned. For most GWASs, the ratio of the number of SNPs and the number of subjects is greater than or close to one, making the sample covariance matrix ill-conditioned ^52, 53^. In these cases, smaller eigenvalues of the sample covariance matrix are underestimated ^52^, leading to inflated false positives in the gene-level association testing. To solve this issue, we choose to adopt the POET estimator ^54^, a principal orthogonal complement thresholding approach, to obtain a well-conditioned covariance matrix via sparse shrinkage under a high-dimensional setting. The estimator of *V* = {*V*_*jl*_; *j, l* = 1, *…, K*} is defined as 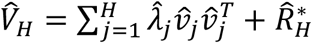, where 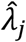 is the *j*th eigenvalues of the covariance matrix with corresponding eigenvector 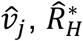 is obtained from applying adaptive thresholding on 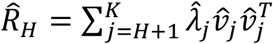, and *H* is the number of spiked eigenvalues. The degree of shrinkage is determined by a tuning parameter, and we choose one so that the positive definiteness of the estimated sparse covariance matrix is guaranteed. Notably, other sparse covariance matrix estimators ^52, 53, 55, 56^ can also be used in a similar fashion.

### Gene-level associations for rare variants

Recent advances in next-generation sequencing technology have made it possible to extend association testing to rare variants, which can explain additional disease risk or trait variability ^33, 35, 57^. Previous work ^58^ has demonstrated that the gene-level testing of rare variants is powerful and able to achieve well-controlled type I error as long as the correlation matrix of single-variant test statistics can be accurately estimated. Here, we recover the burden test statistics from GWAS summary statistics for the gene-level association testing of rare variants. Suppose that a total of *M* rare variants residing in a gene are genotyped. Let *U* = {*U*_*j*_; *j* = 1, *…, M*} and *C* = {*C*_*jl*_; *j, l* = 1, *…, M*} be the score statistic and the corresponding covariance matrix for testing the null hypothesis of no association. Under *H*_0_, the burden test statistic 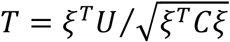 follows a standard normal distribution, where *ξ*_*M×*1_ = (1, ⋯, 1)^*T*^. The GWAS summary statistics do not contain *U* and *C*. We approximate *U*_*j*_ and *C*_*jl*_ by

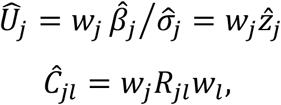

where *R* is the correlation or covariance matrix of 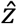 and 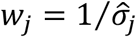 is an empirical approximation to 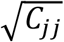. Denote *w* = (*w*_1_, *…, w*_*M*_)^*T*^. The burden test uses 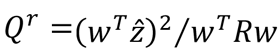, which follows the 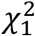 distribution under the null.

### Joint analysis for common and rare variants

Existing methods either remove rare variants from the analysis ^15, 17^ or do not differentiate common and rare variants when only summary statistics are available ^16^. Yet, existing GWASs have successfully uncovered both common and rare variants associated with complex traits and diseases ^23, 33, 35, 57^, and rare variants should therefore not be ignored in the enrichment analysis. To incorporate rare variants into the common-variant gene association testing framework, we collapse genotypes of all rare variants within a gene to construct a pseudo-SNP. We then treat the aggregated pseudo-SNP as a common variant and concatenate the z-scores 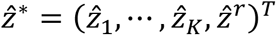, where the first *K* elements are from the common variants and 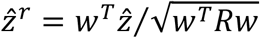 is from the burden test statistic for the combined rare variants. A joint chi-square test for common and rare variants is performed as below:

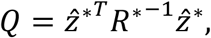

which has the 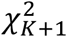 distribution under the null hypothesis. *R*^∗^ can be estimated using POET shrinkage with the pseudo-SNP included.

### Gene-gene correlation

Proximal genes that share *cis*-SNPs inherit LD from SNPs and result in correlations among genes. Since the correlations between genes are caused by LD between SNPs, which quickly drops off as a function of distance, we adopt a sliding-window approach to only compute correlations for pairs of genes within a certain distance from each. It is worth noting that this also significantly reduces the computational burden. Specifically, let *N* be the number of genes from the same chromosome, and we adopt a sliding window of size *d* to estimate the sparse covariance matrix among genes {*G*_1_, *…, G*_*d*_}, {*G*_2_, *…, G*_*d+*1_}, *…*, {*G*_*N−d+*1_, *…, G*_*N*_}, respectively. By default, we set *d* = 10 so that gene-wise correlations can be recovered for a gene with its 18 neighboring genes (see Supplementary Figure S6 for the effect of sliding window size on EPIC’s performance). Similar to MAGMA, correlations are only computed for pairs of genes within 5 megabases by default.

Recall that the gene-level association statistics are chi-square statistics in a quadratic form. Within a specific window, the gene-wise correlations are obtained via transformations of the SNP-wise LD information. Let 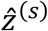 and 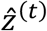 be the SNP-wise z-scores for genes *s* and *t*, respectively. Let 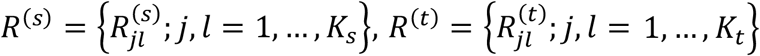, and 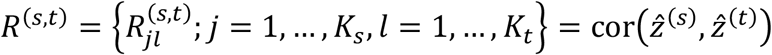 be the within- and between-gene correlation matrices obtained from the POET shrinkage estimation. We take advantage of the Cholesky decomposition to obtain the gene-gene correlation between 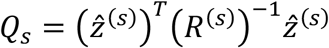 and 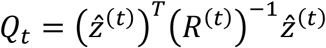:

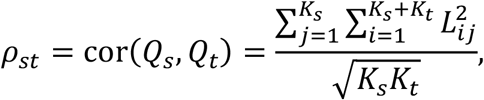

where *L*_*ij*_’s are entries of a lower triangular matrix *L* such that 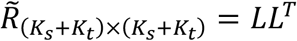 and

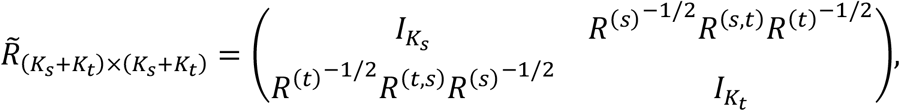

*l*_*K*_ is the identity matrix with dimension *K*. The full derivation is detailed in Supplementary Note S1. When rare variants are included in the framework, gene-gene correlations are calculated similarly by aggregating all rare variants that reside in a gene as a pseudo-SNP.

### Prioritizing trait-relevant cell type(s)

To identify cell-type-specific enrichment for a specific trait of interest, we devise a regression framework based on generalized least squares to identify risk loci enrichment. The key underlying hypothesis is that if a particular cell type influences a trait, more GWAS polygenic signals would be concentrated in genes with greater cell-type-specific gene expression. Under this hypothesis, genes that are significantly associated with lipid traits are expected to be highly expressed in the liver since the liver is known to participate in cholesterol regulation. This relationship between the GWAS association signals and the gene expression specificity is modeled as below.

Let *Q*_*g*_ be the gene-level chi-square association test statistic for gene *g*. To account for the different number of SNPs within each gene, we adjust the degree of freedom of *K*_*g*_ *+* 1 to obtain *Y*_*g*_ = *Q*_*g*_*/*(*K*_*g*_ *+* 1), which is included as the outcome variable. Note that under the null, *Y*_*g*_ has mean of 1 and variance of 2*/*(*K*_*g*_ *+* 1). For each cell type *c*, to test for its enrichment we fit a separate regression using its cell-type-specific gene expression *E*_*cg*_ (reads per kilobase million (RPKM) or transcripts per million (TPM)) as a dependent variable. To account for the baseline gene expression ^24^, we also include another covariate 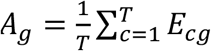, which is the average gene expression across all *T* cell types. Taken together, we have

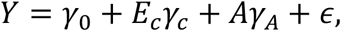

where *ϵ* has a multivariate normal distribution with mean 0 and covariance *σ*^2^*W, W* = *DPD*^*T*^, 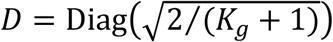, and *P* = {*ρ*_*st*_} is the gene-gene correlation matrix of the chi-square statistics. We adopt the generalized least squares approach to fit the model and perform a one-sided test against the alternative *γ*_*c*_ *>* 0, under which the gene-level association signals positively correlated with the cell-type-specific expression. For a significantly enriched cell type, we further carry out a statistical influence test to identify a set of cell-type-specific influential genes, using the DFBETAS statistics ^59^—large values of DFBETAS indicate observations (i.e., genes) that are influential in estimating *γ*_*c*_. With a size-adjusted cutoff 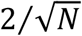, where *N* is the number of genes used in the cell-type-specific enrichment analysis, significantly influential genes allow for further pathway or gene set enrichment analyses.

### GWAS summary statistics and transcriptomic data processing

We adopt GWAS summary statistics of eight traits, including four lipid traits ^31^ (low-density lipoprotein cholesterol (LDL), high-density lipoprotein cholesterol (HDL), total cholesterol (TC), and triglyceride levels (TG)), three neuropsychiatric disorders ^32, 33, 34^ (schizophrenia (SCZ), bipolar disorder (BIP), and schizophrenia and bipolar disorder (SCZBIP)), and type 2 diabetes ^35^ (T2Db). The relevant tissues involved in these traits are well known/studied – liver for the lipid traits, brain for the neuropsychiatric disorders, and pancreas for the T2Db – and we use this as ground truths to demonstrate EPIC and to benchmark against other methods. See Supplementary Table S1 for more information on the GWASs.

For each trait, we obtain SNP-level summary statistics and apply stringent quality control procedures to the data. We restrict our analyses to autosomes, filter out SNPs not in the 1000 Genomes Project Phase 3 reference panel, and remove SNPs with mismatched reference SNP ID numbers. We exclude SNPs from the major histocompatibility complex (MHC) region due to complex LD architecture ^15, 18, 21^. In addition to SNP filtering, we align alleles of each SNP against those of the reference panel to harmonize the effect alleles of all processed GWAS summary statistics. A gene window is defined with 10kb upstream and 1.5kb downstream of each gene ^9^, and SNPs residing in the windows are assigned to the corresponding genes.

In the analysis that follows, we uniformly report results using a minor allele frequency (MAF) cutoff of 1% to define common and rare variants (see Supplementary Figure S7 for enrichment results with different MAF cutoffs). To reduce the computational cost and to alleviate the multicollinearity problem, we perform LD pruning using PLINK ^60^ with a threshold of *r*^2^ *≤* 0.8 to obtain a set of pruned-in common variants, followed by a second-round of LD pruning if the number of common SNPs per gene exceeds 200. See Supplementary Figure S8 for results with varying LD-pruning thresholds. For rare variants, we only carry out a gene-level rare variant association testing if the minor allele count (MAC), defined as the total number of minor alleles across subjects and SNPs within the gene, exceeds 20. We report the number of SNPs (common variants and rare variants), the number of genes, and the number of SNPs per gene for each GWAS trait in Supplementary Table S4.

We adopt a unified framework to process all transcriptomic data. For scRNA-seq data, we follow the Seurat ^61^ pipeline to perform gene- and cell-wise quality controls and focus on the top 8000 highly variable genes. Cell-type-specific RPKMs are calculated by combining read or UMI counts from all cells of a specific cell type, followed by log2 transformation with an added pseudo-count. For tissue-specific bulk RNA-seq data from GTEx, we first calculate a tissue specificity score for each gene ^18, 62^, and only focus on genes that are highly specific in at least one tissue. See Supplementary Note S2 for more details. We then perform log2 transformation on the tissue-specific TPM measurements with an added pseudo-count.

### Benchmarking against RolyPoly, LDSC-SEG, and MAGMA

We benchmarked EPIC against three existing approaches: RolyPoly ^15^, LDSC-SEG ^17^, and MAGMA ^16^. For all methods, we used RPKMs for each cell type and TPMs for each GTEx tissue in the benchmarking analysis. We made gene annotations the same for RolyPoly, MAGMA, and EPIC by defining the gene window as 10kb upstream and 1.5kb downstream of each gene. For LDSC-SEG, as recommended by the authors ^17^, the window size is set to be 100kb up and downstream of each gene’s transcribed region. Since all methods adopt a hypothesis testing framework to identify trait-relevant cell type(s), for each pair of trait and cell type, we reported and compared the corresponding *p*-values from the different methods.

RolyPoly takes as input GWAS summary statistics, gene expression data, gene annotations, and LD matrix from the 1000 Genomes Project Phase 3. As recommended by the developer for RolyPoly ^15^, we scaled the gene expression for each gene across cell types and took the absolute values of the scaled expression values. We performed 100 block bootstrapping iterations to test whether a cell-type-specific gene expression annotation was significantly enriched in a joint model across all cell types. We also benchmarked LDSC-SEG, which computes *t*-statistics to quantify differential expression for each gene across cell types. We annotated genome-wide SNPs using the top 10% genes with the highest positive *t*-statistics and applied stratified LDSC to test the heritability enrichment of the annotations that were attributed to specifically expressed genes for each cell type. For MAGMA, we first obtained gene-level association statistics using MAGMA v1.08. We then carried out the gene-property analysis proposed in Watanabe et al. ^24^, with technical confounders being controlled by default, to test the positive relationship between cell-type specificity of gene expression and genetic associations.

## Discussion

Over the last one and half decades, GWASs have successfully identified and replicated genetic variants associated with various complex traits. Meanwhile, bulk-tissue and single-cell transcriptomic sequencing allow tissue- and cell-type-specific gene expression characterization and have seen rapid technological development with ever-increasing sequencing capacities and throughputs. Here, we propose EPIC to address the problem of how GWAS summary statistics should be integrated with scRNA-seq data to prioritize trait-relevant cell type(s) and to elucidate disease etiology. To our best knowledge, EPIC is the first method that prioritizes cell type(s) for both common and rare variants with a rigorous statistical framework that properly accounts for both within- and between-gene correlations. We demonstrate EPIC’s effectiveness and outperformance compared to existing methods with extensive benchmark and validation studies.

For scRNA-seq data, all existing methods, including EPIC, resort to pre-clustered/annotated cell types and average across cells to obtain cell-type-specific expression profiles. However, scRNA-seq goes beyond the mean measurements ^63, 64^, and how to make the best use of gene expression dispersion, nonzero fraction, and other aspects of its distribution needs further method development ^65^. Additionally, while many efforts have been devoted to identifying enrichment of discretized cell types, how to carry out enrichment analysis for transient cell states needs further investigation. Last but not least, when multiple scRNA-seq datasets are available across different experiments, protocols, or species, borrowing information from additional sources can potentially boost the performance and increase the robustness of the enrichment analysis ^66^. While it is nontrivial to directly perform gene expression data integration, a cross-dataset conditional analysis workflow was proposed by Watanabe et al. ^24^ to evaluate the association of cell types based on multiple independent scRNA-seq datasets. However, the linear conditional analysis may not be sufficient to capture any nonlinear batch effects ^61, 67^.

It is also worth noting that CoCoNet, MAGMA, and EPIC first carry out a gene-level association test so that the summary statistics and expressions are unified to be gene-specific. They integrate SNP-wise summary statistics in different ways, yet for all methods, SNPs need to be first annotated to genes based on a window surrounding each gene. While RolyPoly and LDSC-SEG model on the SNP level directly, each SNP still needs to be assigned to a gene so that the gene expression can be used as an SNP annotation. There is not a consensus on how to most accurately assign SNPs to genes, and more importantly, one would only be able to perform gene annotations for SNPs that reside in gene bodies or promoter regions. Meanwhile, a large number of GWAS hits are in the non-coding regions, and their functions are yet to be fully understood. EPIC’s framework can be easily extended to infer enrichment of non-coding variants when combined with the single-cell assay for transposase-accessible chromatin using sequencing (ATAC-seq) data ^68, 69^. Additionally, cell-type-specific expression quantitative trait loci from the non-coding regions ^70^ can also be integrated with the second-step gene-property analysis to boost power and to infer enrichment of non-coding variants.

## Data Availability

GWAS summary statistics are downloaded from public repositories listed in Supplementary Table S1. Genotypes from the 1000 Genomes Project reference panel are available at https://ctg.cncr.nl/software/magma. Bulk RNA-seq and scRNA-seq data are downloaded from GTEx v8 at http://www.gtexportal.org. ScRNA-seq read counts from two pancreatic islet studies are publicly available with accession GSE81433 ^6^ and E-MTAB-5061 ^8^. We obtain a list of human housekeeping genes from the Housekeeping and Reference Transcript Atlas ^37^ at https://housekeeping.unicamp.br.

## Code Availability

EPIC is compiled as an open-source R package available at https://github.com/rujinwang/EPIC.

## Acknowledgments

This work was supported by the National Institutes of Health (NIH) grant P01 CA142538 (to D.L. and Y.J.), R35 GM138342 (to Y.J.), and R01 HG009974 (to D.L.). The authors thank Drs. Yun Li, Michael Love, Karen Mohlke, and Jason Stein for helpful discussions and comments, and Drs. Alkes Price, Diego Calderon, and Kyoko Watanabe for providing support and insight on existing methods.

## Authors’ Contributions

Y.J. and D.L. initiated and envisioned the study. R.W., D.L., and Y.J. formulated the model; R.W. developed and implemented the algorithm. R.W., D.L., and Y.J. performed data analysis. R.W. and Y.J. wrote the manuscript, which was edited by D.L..

## Competing Interests

The authors declare no competing interests.

## Notes

### Competing Interest Statement

The authors have declared no competing interest.

## References

1. Lang UE, Puls I, Muller DJ, Strutz-Seebohm N, Gallinat J. Molecular mechanisms of schizophrenia. Cell Physiol Biochem 20, 687–702 (2007).

2. Ongen H, et al. Estimating the causal tissues for complex traits and diseases. Nat Genet 49, 1676–1683 (2017).

3. Raj T, et al. Polarization of the effects of autoimmune and neurodegenerative risk alleles in leukocytes. Science 344, 519–523 (2014).

4. Uhlhaas PJ, Singer W. Abnormal neural oscillations and synchrony in schizophrenia. Nat Rev Neurosci 11, 100–113 (2010).

5. Xiao X, Chang H, Li M. Molecular mechanisms underlying noncoding risk variations in psychiatric genetic studies. Mol Psychiatry 22, 497–511 (2017).

6. Baron M, et al. A Single-Cell Transcriptomic Map of the Human and Mouse Pancreas Reveals Inter- and Intra-cell Population Structure. Cell Syst 3, 346–360 e344 (2016).

7. Habib N, et al. Massively parallel single-nucleus RNA-seq with DroNc-seq. Nat Methods 14, 955–958 (2017).

8. Segerstolpe A, et al. Single-Cell Transcriptome Profiling of Human Pancreatic Islets in Health and Type 2 Diabetes. Cell Metab 24, 593–607 (2016).

9. Skene NG, et al. Genetic identification of brain cell types underlying schizophrenia. Nat Genet 50, 825–833 (2018).

10. Barbeira AN, et al. Exploring the phenotypic consequences of tissue specific gene expression variation inferred from GWAS summary statistics. Nat Commun 9, 1825 (2018).

11. Bryois J, et al. Genetic identification of cell types underlying brain complex traits yields insights into the etiology of Parkinson’s disease. Nature Genetics, 1–12 (2020).

12. Lake BB, et al. Integrative single-cell analysis of transcriptional and epigenetic states in the human adult brain. Nat Biotechnol 36, 70–80 (2018).

13. Sudlow C, et al. UK biobank: an open access resource for identifying the causes of a wide range of complex diseases of middle and old age. PLoS Med 12, e1001779 (2015).

14. Muus C, et al. Integrated analyses of single-cell atlases reveal age, gender, and smoking status associations with cell type-specific expression of mediators of SARS-CoV-2 viral entry and highlights inflammatory programs in putative target cells. bioRxiv, 2020.2004.2019.049254 (2020).

15. Calderon D, et al. Inferring Relevant Cell Types for Complex Traits by Using Single-Cell Gene Expression. Am J Hum Genet 101, 686–699 (2017).

16. de Leeuw CA, Mooij JM, Heskes T, Posthuma D. MAGMA: generalized gene-set analysis of GWAS data. PLoS Comput Biol 11, e1004219 (2015).

17. Finucane HK, et al. Heritability enrichment of specifically expressed genes identifies disease-relevant tissues and cell types. Nat Genet 50, 621–629 (2018).

18. Shang L, Smith JA, Zhou X. Leveraging gene co-expression patterns to infer trait-relevant tissues in genome-wide association studies. PLoS Genet 16, e1008734 (2020).

19. Zhu H, Shang L, Zhou X. A Review of Statistical Methods for Identifying Trait-Relevant Tissues and Cell Types. Front Genet 11, 587887 (2020).

20. Bulik-Sullivan BK, et al. LD Score regression distinguishes confounding from polygenicity in genome-wide association studies. Nat Genet 47, 291–295 (2015).

21. Finucane HK, et al. Partitioning heritability by functional annotation using genome-wide association summary statistics. Nat Genet 47, 1228–1235 (2015).

22. Jagadeesh KA, et al. Identifying disease-critical cell types and cellular processes across the human body by integration of single-cell profiles and human genetics. 2021.2003.2019.436212 (2021).

23. Bryois J, et al. Genetic identification of cell types underlying brain complex traits yields insights into the etiology of Parkinson’s disease. Nat Genet 52, 482–493 (2020).

24. Watanabe K, Umicevic Mirkov M, de Leeuw CA, van den Heuvel MP, Posthuma D. Genetic mapping of cell type specificity for complex traits. Nat Commun 10, 3222 (2019).

25. Kalra G, et al. Biological insights from multi-omic analysis of 31 genomic risk loci for adult hearing difficulty. PLoS Genet 16, e1009025 (2020).

26. Timshel PN, Thompson JJ, Pers TH. Genetic mapping of etiologic brain cell types for obesity. Elife 9, (2020).

27. Tran MN, et al. Single-nucleus transcriptome analysis reveals cell type-specific molecular signatures across reward circuitry in the human brain. bioRxiv, x2020.2010.2007.329839 (2020).

28. Yurko R, Roeder K, Devlin B, G’Sell M. H-MAGMA, inheriting a shaky statistical foundation, yields excess false positives. Ann Hum Genet 85, 97–100 (2021).

29. Wu MC, Lee S, Cai T, Li Y, Boehnke M, Lin X. Rare-variant association testing for sequencing data with the sequence kernel association test. Am J Hum Genet 89, 82–93 (2011).

30. Lin DY, Tang ZZ. A general framework for detecting disease associations with rare variants in sequencing studies. Am J Hum Genet 89, 354–367 (2011).

31. Willer CJ, et al. Discovery and refinement of loci associated with lipid levels. Nat Genet 45, 1274–1283 (2013).

32. Bipolar D, Schizophrenia Working Group of the Psychiatric Genomics Consortium. Electronic address drve, Bipolar D, Schizophrenia Working Group of the Psychiatric Genomics C. Genomic Dissection of Bipolar Disorder and Schizophrenia, Including 28 Subphenotypes. Cell 173, 1705–1715 e1716 (2018).

33. Schizophrenia Working Group of the Psychiatric Genomics C. Biological insights from 108 schizophrenia-associated genetic loci. Nature 511, 421–427 (2014).

34. Stahl EA, et al. Genome-wide association study identifies 30 loci associated with bipolar disorder. Nat Genet 51, 793–803 (2019).

35. Mahajan A, et al. Fine-mapping type 2 diabetes loci to single-variant resolution using high-density imputation and islet-specific epigenome maps. Nat Genet 50, 1505–1513 (2018).

36. Consortium GT. The GTEx Consortium atlas of genetic regulatory effects across human tissues. Science 369, 1318–1330 (2020).

37. Hounkpe BW, Chenou F, de Lima F, De Paula EV. HRT Atlas v1.0 database: redefining human and mouse housekeeping genes and candidate reference transcripts by mining massive RNA-seq datasets. Nucleic Acids Res 49, D947–D955 (2021).

38. Ko CW, Qu J, Black DD, Tso P. Regulation of intestinal lipid metabolism: current concepts and relevance to disease. Nat Rev Gastroenterol Hepatol 17, 169–183 (2020).

39. Field FJ, Kam NT, Mathur SN. Regulation of cholesterol metabolism in the intestine. Gastroenterology 99, 539–551 (1990).

40. Severson DL. Regulation of lipid metabolism in adipose tissue and heart. Can J Physiol Pharmacol 57, 923–937 (1979).

41. Coppack SW, Patel JN, Lawrence VJ. Nutritional regulation of lipid metabolism in human adipose tissue. Exp Clin Endocrinol Diabetes 109 Suppl 2, S202–214 (2001).

42. Cerf ME. Beta cell dysfunction and insulin resistance. Front Endocrinol (Lausanne) 4, 37 (2013).

43. Donath MY, et al. Mechanisms of beta-cell death in type 2 diabetes. Diabetes 54 Suppl 2, S108–113 (2005).

44. Chandra R, Liddle RA. Neural and hormonal regulation of pancreatic secretion. Curr Opin Gastroenterol 25, 441–446 (2009).

45. Chandra R, Liddle RA. Recent advances in the regulation of pancreatic secretion. Curr Opin Gastroenterol 30, 490–494 (2014).

46. Washabau RJ. Chapter 1 - Integration of Gastrointestinal Function. In: Canine and Feline Gastroenterology (ed^(eds Washabau RJ, Day MJ). W.B. Saunders (2013).

47. Huang da W, Sherman BT, Lempicki RA. Systematic and integrative analysis of large gene lists using DAVID bioinformatics resources. Nat Protoc 4, 44–57 (2009).

48. Huang da W, Sherman BT, Lempicki RA. Bioinformatics enrichment tools: paths toward the comprehensive functional analysis of large gene lists. Nucleic Acids Res 37, 1–13 (2009).

49. Pardinas AF, et al. Common schizophrenia alleles are enriched in mutation-intolerant genes and in regions under strong background selection. Nat Genet 50, 381–389 (2018).

50. Fromer M, et al. Gene expression elucidates functional impact of polygenic risk for schizophrenia. Nat Neurosci 19, 1442–1453 (2016).

51. Genomes Project C, et al. An integrated map of genetic variation from 1,092 human genomes. Nature 491, 56–65 (2012).

52. Ledoit O, Wolf M. A well-conditioned estimator for large-dimensional covariance matrices. Journal of Multivariate Analysis 88, 365–411 (2004).

53. Cai T, Liu W. Adaptive Thresholding for Sparse Covariance Matrix Estimation. Journal of the American Statistical Association 106, 672–684 (2011).

54. Fan J, Liao Y, Mincheva M. Large Covariance Estimation by Thresholding Principal Orthogonal Complements. J R Stat Soc Series B Stat Methodol 75, (2013).

55. Bickel PJ, Levina E. Covariance Regularization by Thresholding. Ann Stat 36, 2577–2604 (2008).

56. Ledoit O, Wolf M. Spectrum estimation: A unified framework for covariance matrix estimation and PCA in large dimensions. Journal of Multivariate Analysis 139, 360–384 (2015).

57. Lange LA, et al. Whole-exome sequencing identifies rare and low-frequency coding variants associated with LDL cholesterol. Am J Hum Genet 94, 233–245 (2014).

58. Hu YJ, et al. Meta-analysis of gene-level associations for rare variants based on single-variant statistics. Am J Hum Genet 93, 236–248 (2013).

59. Belsley DA, Kuh E, Welsch RE. Regression diagnostics : identifying influential data and sources of collinearity. Wiley (1980).

60. Chang CC, Chow CC, Tellier LC, Vattikuti S, Purcell SM, Lee JJ. Second-generation PLINK: rising to the challenge of larger and richer datasets. Gigascience 4, 7 (2015).

61. Stuart T, et al. Comprehensive Integration of Single-Cell Data. Cell 177, 1888–1902 e1821 (2019).

62. Sonawane AR, et al. Understanding Tissue-Specific Gene Regulation. Cell Rep 21, 1077–1088 (2017).

63. Jiang Y, Zhang NR, Li M. SCALE: modeling allele-specific gene expression by single-cell RNA sequencing. Genome Biol 18, 74 (2017).

64. Korthauer KD, et al. A statistical approach for identifying differential distributions in single-cell RNA-seq experiments. Genome Biol 17, 222 (2016).

65. Wang J, et al. Gene expression distribution deconvolution in single-cell RNA sequencing. Proc Natl Acad Sci U S A 115, E6437–E6446 (2018).

66. Dong M, et al. SCDC: bulk gene expression deconvolution by multiple single-cell RNA sequencing references. Brief Bioinform 22, 416–427 (2021).

67. Haghverdi L, Lun ATL, Morgan MD, Marioni JC. Batch effects in single-cell RNA-sequencing data are corrected by matching mutual nearest neighbors. Nat Biotechnol 36, 421–427 (2018).

68. Urrutia E, Chen L, Zhou H, Jiang Y. Destin: toolkit for single-cell analysis of chromatin accessibility. Bioinformatics 35, 3818–3820 (2019).

69. Granja JM, et al. ArchR is a scalable software package for integrative single-cell chromatin accessibility analysis. Nat Genet 53, 403–411 (2021).

70. van der Wijst MGP, et al. Single-cell RNA sequencing identifies celltype-specific cis-eQTLs and co-expression QTLs. Nat Genet 50, 493–497 (2018).

