## Supplement for "EPIC: inferring relevant cell types for complex traits by integrating genome-wide association studies and single-cell RNA sequencing"

### Supplementary Note

#### Note S1: Derivation of gene-gene correlations

We first consider a special case that each gene only contains one SNP. Let $X \sim N(0,1)$, $Y \sim N(0,1)$ and $\mathrm{Cov}(X,Y) = \mathrm{Cor}(X,Y) = \rho$. The goal is to compute $\mathrm{Cov}(X^2, Y^2)$ and $\mathrm{Cor}(X^2, Y^2)$.

Let $X = U_1$, $Y = \rho U_1 + \sqrt{1-\rho^2}U_2$, where $U_1, U_2 \overset{iid}{\sim} N(0,1)$.

$$\begin{aligned}\mathrm{Cov}(X^2, Y^2) &= \mathrm{Cov}\left(U_1^2, \left(\rho U_1 + \sqrt{1-\rho^2}U_2\right)^2\right)\\ &= \rho^2\mathrm{Cov}(U_1^2, U_1^2) + 2\rho\sqrt{1-\rho^2}\mathrm{Cov}(U_1^2, U_1U_2) + (1-\rho^2)\mathrm{Cov}(U_1^2, U_2^2)\\ &= \rho^2\mathrm{Cov}(U_1^2, U_1^2) = 2\rho^2,\end{aligned}$$

by making use of $\mathrm{Cov}(U_1, U_1) = 1$, $\mathrm{Cov}(U_1, U_2) = 0$, and $\mathrm{Cov}(U_1^2, U_1U_2) = 0$.

Now we consider the generalized case. Let $X \sim \mathrm{MVN}(0, I_p)$, $Y \sim \mathrm{MVN}(0, I_q)$, and $\mathrm{Cov}\binom{X}{Y} = \mathrm{Cor}\binom{X}{Y} = R_{(p+q)\times(p+q)} = \begin{pmatrix} I_p & R_{XY} \\ R_{XY}^T & I_q \end{pmatrix}$. Suppose $U \sim \mathrm{MVN}(0, I_{p+q})$. We take advantage of Cholesky decomposition:

$$R = LL^T,$$

where

$$L = \begin{pmatrix} L_{11} & 0 & 0 & \cdots & 0 \\ L_{21} & L_{22} & 0 & \cdots & 0 \\ L_{31} & L_{32} & L_{33} & \cdots & 0 \\ \vdots & \vdots & \vdots & \vdots & \vdots \\ L_{(p+q)1} & L_{(p+q)2} & L_{(p+q)3} & \cdots & L_{(p+q)(p+q)} \end{pmatrix}.$$

Therefore, $\binom{X}{Y} = LU$. For each $i, j, k = 1, \cdots, (p+q)$ and $i \neq j \neq k$, we have $\mathrm{Var}(U_i) = 2$, $\mathrm{Cov}(U_i^2, U_j^2) = 0$, $\mathrm{Cov}(U_i^2, U_iU_j) = 0$, $\mathrm{Cov}(U_iU_j, U_iU_j) = 1$, and $\mathrm{Cov}(U_iU_j, U_iU_k) = 0$.

$$\begin{aligned}\mathrm{Cov}(X^TX, Y^TY) &= \mathrm{Cov}\left(\left(L_{[1:p,\cdot]}U\right)^T\left(L_{[1:p,\cdot]}U\right), \left(L_{[(p+1):(p+q),\cdot]}U\right)^T\left(L_{[(p+1):(p+q),\cdot]}U\right)\right)\\ &= \left(\sum_{i=1}^{p} L_{i1}^2\right)\mathrm{Var}(U_1^2)\left(\sum_{i=p+1}^{p+q} L_{i1}^2\right) + \cdots + \left(\sum_{i=1}^{p} L_{ip}^2\right)\mathrm{Var}\left(U_p^2\right)\left(\sum_{i=p+1}^{p+q} L_{ip}^2\right)\\ &+ 4\sum_{m=1}^{p-1}\sum_{n=m+1}^{p}\mathrm{Cov}(U_mU_n, U_mU_n)\left(\sum_{i=1}^{p} L_{im}L_{in}\right)\left(\sum_{i=p+1}^{p+q} L_{im}L_{in}\right)\\ &= 2\left(\left(\sum_{i=1}^{p} L_{i1}^2\right)\left(\sum_{i=p+1}^{p+q} L_{i1}^2\right) + \cdots + \left(\sum_{i=1}^{p} L_{ip}^2\right)\left(\sum_{i=p+1}^{p+q} L_{ip}^2\right)\right)\\ &+ 4\sum_{m=1}^{p-1}\sum_{n=m+1}^{p}\left(\left(\sum_{i=1}^{p} L_{im}L_{in}\right)\left(\sum_{i=p+1}^{p+q} L_{im}L_{in}\right)\right),\end{aligned}$$

where $L_{[1:p,\cdot]}$ indicates the submatrix of $L$ from $i$th to $j$th row.

Specifically, given $R = \begin{pmatrix} I_p & R_{XY} \\ R_{XY}^T & I_q \end{pmatrix}$ and the Cholesky decomposition steps, $L_{[1:p,1:p]} = I_p$.

As a result,

$$\mathrm{Cov}(X^T X, Y^T Y) = 2\left(\left(\sum_{i=p+1}^{p+q} L_{i1}^2\right) + \cdots + \left(\sum_{i=p+1}^{p+q} L_{ip}^2\right)\right)$$

For the chi-square gene-level association statistics, let $X = {R^{(s)}}^{-1/2}\hat{z}^{(s)}$, $Y = {R^{(t)}}^{-1/2}\hat{z}^{(t)}$, so that $X \sim \mathrm{MVN}(0, I_{K_s})$, $Y \sim \mathrm{MVN}(0, I_{K_t})$, and

$$\mathrm{Cov}\begin{pmatrix} X \\ Y \end{pmatrix} = \mathrm{Cor}\begin{pmatrix} X \\ Y \end{pmatrix} = \tilde{R}_{(K_s+K_t)\times(K_s+K_t)} = \begin{pmatrix} I_{K_s} & {R^{(s)}}^{-1/2} R^{(s,t)} {R^{(t)}}^{-1/2} \\ {R^{(t)}}^{-1/2} R^{(t,s)} {R^{(s)}}^{-1/2} & I_{K_t} \end{pmatrix}$$

Denote $Q_s = \left(\hat{z}^{(s)}\right)^T \left(R^{(s)}\right)^{-1} \hat{z}^{(s)}$ and $Q_t = \left(\hat{z}^{(t)}\right)^T \left(R^{(t)}\right)^{-1} \hat{z}^{(t)}$. We perform Cholesky decomposition on $\tilde{R} = LL^T$ and obtain

$$\mathrm{cov}(Q_s, Q_t) = 2\left(\sum_{j=1}^{K_s} \sum_{i=1}^{K_s+K_t} L_{ij}^2\right)$$

As a result,

$$\rho_{st} = \mathrm{cor}(Q_s, Q_t) = \frac{\sum_{j=1}^{K_s} \sum_{i=1}^{K_s+K_t} L_{ij}^2}{\sqrt{K_s K_t}},$$

where $L_{ij}$'s are entries of a lower triangular matrix $L$ such that $\tilde{R}_{(K_s+K_t)\times(K_s+K_t)} = LL^T$.

**Note S2: Tissue-specific gene selection in bulk GTEx dataset**

We compute gene specificity score for gene $i$ and tissue $t$ as follows:

$$s_i^t = \frac{\mathrm{median}(e_i^t) - \mathrm{median}\left(e_i^{all}\right)}{\mathrm{IQR}\left(e_i^{all}\right)},$$

where $\mathrm{median}(e_i^t)$ is the median expression of gene $i$ in a particular tissue $t$; $\mathrm{median}\left(e_i^{all}\right)$ and $\mathrm{IQR}\left(e_i^{all}\right)$ are the median and IQR of its expression across all samples. We define genes with a gene specificity score $s_i^t \geq 5$ in any tissue $t$ as tissue-specific genes, which are selected in subsequent analyses.

### Supplementary Tables

#### Table S1: Summary of GWAS studies and transcriptomic studies.

**(A)** Summary information for eight GWAS studies

| Phenotype | Abbreviation | # of subjects | Reference | URL |
|---|---|---|---|---|
| Low-density lipoprotein cholesterol | LDL | 188,577 | Willer et al., 2013 [1] | http://csg.sph.umich.edu/willer/public/lipids2013/ |
| High-density lipoprotein cholesterol | HDL | 188,577 | Willer et al., 2013 [1] | http://csg.sph.umich.edu/willer/public/lipids2013/ |
| Total cholesterol | TC | 188,577 | Willer et al., 2013 [1] | http://csg.sph.umich.edu/willer/public/lipids2013/ |
| Triglyceride levels | TG | 188,577 | Willer et al., 2013 [1] | http://csg.sph.umich.edu/willer/public/lipids2013/ |
| Schizophrenia | SCZ | 79,845 | Ripke et al., 2014 [2] | https://www.med.unc.edu/pgc/download-results/ |
| Schizophrenia | SCZ2 | 105,318 | Pardinas et al., 2018 [3] | https://walters.psycm.cf.ac.uk/ |
| Bipolar disorder | BIP | 51,710 | Stahl et al., 2019 [4] | https://www.med.unc.edu/pgc/download-results/ |
| Schizophrenia and bipolar disorder | SCZBIP | 107,620 | Ruderfer et al., 2018 [5] | https://www.med.unc.edu/pgc/download-results/ |
| Type 2 diabetes | T2Db | 898,130 | Mahajan et al., 2018b [6] | http://diagram-consortium.org/downloads.html |

**(B)** Summary information for transcriptomic studies

| Name | Tissue or cell type | Technology | Reference |
|---|---|---|---|
| GTEx bulk | 45 tissues from 980 donors | RNA-seq | GTEx Consortium, 2020 [7] |
| Pancreatic islet scRNA-seq: Baron | 13 cell types from 3 healthy donors | InDrop | Baron et al. [8] |
| Pancreatic islet scRNA-seq: Segerstolpe | 12 cell types from 6 healthy donors | SMART-Seq2 | Segerstolpe et al. [9] |
| GTEx scRNA-seq | 10 brain cell types from 5 donors | DroNc-seq | Habib et al. [10] |

**Table S2. Gene-level association testing results of eight GWAS traits.** A final set of 8,708 genes are retained. We selected a list of risk genes within implicated genome-wide significant loci that were reported in the original GWAS [1, 2, 4, 5, 6] for each trait. Significant gene-level associations were detected between all lipid traits and variants in *APOB*, *APOE*, and *CETP*. Meanwhile, *PCSK9*, *ABCG5,* and *ABCG8* exhibited significant associations with LDL and TC. For neuropsychiatric disorders, we examined genes that are relevant to the etiology of schizophrenia, including genes that are targets of therapeutic drugs (*DRD2* and *GRM3*), genes that participate in neuronal calcium signaling (*CACNA1I*), and genes that are involved in synaptic function (*CNTN4* and *SNAP91*) and other neuronal pathways (*FXR1*, *CHRNA3*, *CHRNB4*, and *HCN1*). EPIC's chi-square test approach demonstrates higher power than MAGMA.

(A) LDL: gene-level association testing

| hgID | chr | EPIC | | | MAGMA |
|---|---|---|---|---|---|
| | | Joint | Common | Rare | |
| *PCSK9* | 1 | $1.00 \times 10^{-300}$ | $1.00 \times 10^{-300}$ | $4.61 \times 10^{-2}$ | $5.00 \times 10^{-10}$ |
| *ANGPTL3* | 1 | $2.20 \times 10^{-06}$ | $3.38 \times 10^{-06}$ | $6.00 \times 10^{-2}$ | $3.72 \times 10^{-15}$ |
| *APOB* | 2 | $1.00 \times 10^{-300}$ | $1.00 \times 10^{-300}$ | $3.37 \times 10^{-05}$ | $1.30 \times 10^{-14}$ |
| *ABCG8* | 2 | $8.81 \times 10^{-111}$ | $8.81 \times 10^{-111}$ | $NA$ | $1.15 \times 10^{-12}$ |
| *ABCG5* | 2 | $2.36 \times 10^{-104}$ | $2.36 \times 10^{-104}$ | $NA$ | $5.00 \times 10^{-10}$ |
| *EHBP1* | 2 | $1.08 \times 10^{-09}$ | $1.06 \times 10^{-09}$ | $NA$ | $2.48 \times 10^{-06}$ |
| *TIMD4* | 5 | $3.23 \times 10^{-22}$ | $2.16 \times 10^{-21}$ | $3.89 \times 10^{-2}$ | $3.72 \times 10^{-15}$ |
| *LPA* | 6 | $3.38 \times 10^{-15}$ | $1.57 \times 10^{-15}$ | $6.77 \times 10^{-4}$ | $2.15 \times 10^{-11}$ |
| *NPC1L1* | 7 | $5.16 \times 10^{-29}$ | $2.10 \times 10^{-28}$ | $5.65 \times 10^{-06}$ | $2.07 \times 10^{-12}$ |
| *DNAH11* | 7 | $1.21 \times 10^{-4}$ | $1.17 \times 10^{-4}$ | $NA$ | $3.60 \times 10^{-07}$ |
| *TRIB1* | 8 | $4.89 \times 10^{-24}$ | $1.84 \times 10^{-24}$ | $2.72 \times 10^{-1}$ | $4.24 \times 10^{-13}$ |
| *ABO* | 9 | $1.64 \times 10^{-102}$ | $2.22 \times 10^{-92}$ | $9.59 \times 10^{-05}$ | $5.00 \times 10^{-10}$ |
| *VLDLR* | 9 | $1.16 \times 10^{-08}$ | $1.07 \times 10^{-08}$ | $NA$ | $1.38 \times 10^{-2}$ |
| *GPAM* | 10 | $6.76 \times 10^{-08}$ | $6.76 \times 10^{-08}$ | $NA$ | $1.12 \times 10^{-09}$ |
| *ST3GAL4* | 11 | $7.72 \times 10^{-46}$ | $5.91 \times 10^{-44}$ | $5.41 \times 10^{-1}$ | $7.22 \times 10^{-15}$ |
| *HNF1A* | 12 | $5.57 \times 10^{-21}$ | $5.29 \times 10^{-21}$ | $NA$ | $5.00 \times 10^{-10}$ |
| *HPR* | 16 | $4.41 \times 10^{-38}$ | $2.01 \times 10^{-38}$ | $9.43 \times 10^{-1}$ | $3.46 \times 10^{-11}$ |
| *CETP* | 16 | $9.29 \times 10^{-35}$ | $1.13 \times 10^{-36}$ | $8.77 \times 10^{-16}$ | $6.16 \times 10^{-15}$ |
| *APOE* | 19 | $1.00 \times 10^{-300}$ | $1.00 \times 10^{-300}$ | $4.59 \times 10^{-2}$ | $1.17 \times 10^{-15}$ |

(B) HDL: gene-level association testing

| hgID | chr | EPIC | | | MAGMA |
|---|---|---|---|---|---|
| | | Joint | Common | Rare | |
| *CPS1* | 2 | $2.55 \times 10^{-11}$ | $1.30 \times 10^{-10}$ | $NA$ | $5.77 \times 10^{-3}$ |
| *SLC39A8* | 4 | $8.57 \times 10^{-10}$ | $1.53 \times 10^{-06}$ | $2.90 \times 10^{-1}$ | $1.70 \times 10^{-5}$ |
| *RSPO3* | 6 | $1.81 \times 10^{-10}$ | $1.58 \times 10^{-10}$ | $7.90 \times 10^{-2}$ | $2.01 \times 10^{-08}$ |
| *MLXIPL* | 7 | $8.72 \times 10^{-15}$ | $8.65 \times 10^{-12}$ | $8.84 \times 10^{-1}$ | $1.53 \times 10^{-07}$ |
| *LPL* | 8 | $1.00 \times 10^{-300}$ | $1.00 \times 10^{-300}$ | $1.38 \times 10^{-2}$ | $5.00 \times 10^{-10}$ |
| *TRIB1* | 8 | $8.49 \times 10^{-11}$ | $4.94 \times 10^{-11}$ | $3.81 \times 10^{-1}$ | $8.22 \times 10^{-6}$ |
| *SCARB1* | 12 | $5.25 \times 10^{-74}$ | $4.09 \times 10^{-74}$ | $NA$ | $3.77 \times 10^{-14}$ |
| *LIPC* | 15 | $1.00 \times 10^{-300}$ | $1.00 \times 10^{-300}$ | $5.76 \times 10^{-09}$ | $5.00 \times 10^{-10}$ |
| *CETP* | 16 | $1.00 \times 10^{-300}$ | $1.00 \times 10^{-300}$ | $5.94 \times 10^{-108}$ | $5.00 \times 10^{-10}$ |
| *ABCA8* | 17 | $1.39 \times 10^{-11}$ | $1.11 \times 10^{-15}$ | $4.87 \times 10^{-06}$ | $5.54 \times 10^{-10}$ |
| *LIPG* | 18 | $7.84 \times 10^{-36}$ | $8.16 \times 10^{-36}$ | $NA$ | $3.61 \times 10^{-15}$ |
| *MC4R* | 18 | $4.04 \times 10^{-08}$ | $1.16 \times 10^{-06}$ | $1.74 \times 10^{-3}$ | $7.16 \times 10^{-3}$ |
| *APOE* | 19 | $1.60 \times 10^{-54}$ | $3.25 \times 10^{-55}$ | $7.87 \times 10^{-1}$ | $1.05 \times 10^{-10}$ |
| *HNF4A* | 20 | $5.75 \times 10^{-39}$ | $1.08 \times 10^{-39}$ | $2.59 \times 10^{-1}$ | $3.98 \times 10^{-4}$ |

(C) TC: gene-level association testing

| hgID | chr | EPIC | | | MAGMA |
|---|---|---|---|---|---|
| | | Joint | Common | Rare | |
| *PCSK9* | 1 | $1.72 \times 10^{-237}$ | $7.14 \times 10^{-232}$ | $2.42 \times 10^{-1}$ | $3.24 \times 10^{-13}$ |
| *ANGPTL3* | 1 | $7.74 \times 10^{-16}$ | $1.65 \times 10^{-16}$ | $5.59 \times 10^{-1}$ | $2.78 \times 10^{-16}$ |
| *APOB* | 2 | $1.00 \times 10^{-300}$ | $1.00 \times 10^{-300}$ | $8.45 \times 10^{-4}$ | $1.25 \times 10^{-13}$ |
| *ABCG8* | 2 | $2.67 \times 10^{-114}$ | $1.26 \times 10^{-114}$ | $NA$ | $5.00 \times 10^{-10}$ |
| *ABCG5* | 2 | $4.56 \times 10^{-103}$ | $2.92 \times 10^{-103}$ | $NA$ | $9.55 \times 10^{-13}$ |
| *GCKR* | 2 | $2.21 \times 10^{-60}$ | $1.44 \times 10^{-62}$ | $7.9 \times 10^{-2}$ | $1.74 \times 10^{-14}$ |
| *ABCB11* | 2 | $3.41 \times 10^{-06}$ | $1.26 \times 10^{-05}$ | $1.75 \times 10^{-4}$ | $1.06 \times 10^{-3}$ |
| *UGT1A1* | 2 | $3.86 \times 10^{-4}$ | $7.45 \times 10^{-4}$ | $1.96 \times 10^{-3}$ | $1.12 \times 10^{-06}$ |
| *TIMD4* | 5 | $5.81 \times 10^{-29}$ | $6.27 \times 10^{-27}$ | $2.61 \times 10^{-3}$ | $5.00 \times 10^{-10}$ |
| *LPA* | 6 | $3.01 \times 10^{-14}$ | $9.68 \times 10^{-15}$ | $5.71 \times 10^{-3}$ | $1.62 \times 10^{-09}$ |
| *NPC1L1* | 7 | $1.36 \times 10^{-26}$ | $2.71 \times 10^{-26}$ | $1.76 \times 10^{-06}$ | $2.69 \times 10^{-12}$ |
| *DNAH11* | 7 | $1.23 \times 10^{-4}$ | $1.25 \times 10^{-4}$ | $NA$ | $6.20 \times 10^{-09}$ |
| *TRIB1* | 8 | $3.36 \times 10^{-28}$ | $9.49 \times 10^{-29}$ | $4.59 \times 10^{-1}$ | $1.50 \times 10^{-15}$ |
| *CYP7A1* | 8 | $1.58 \times 10^{-4}$ | $1.55 \times 10^{-4}$ | $NA$ | $1.92 \times 10^{-08}$ |
| *ABO* | 9 | $5.24 \times 10^{-60}$ | $1.12 \times 10^{-59}$ | $6.31 \times 10^{-4}$ | $3.12 \times 10^{-11}$ |
| *VLDLR* | 9 | $1.44 \times 10^{-06}$ | $1.18 \times 10^{-06}$ | $NA$ | $4.49 \times 10^{-2}$ |
| *GPAM* | 10 | $8.68 \times 10^{-10}$ | $8.68 \times 10^{-10}$ | $NA$ | $1.02 \times 10^{-10}$ |
| *ST3GAL4* | 11 | $6.92 \times 10^{-19}$ | $5.13 \times 10^{-18}$ | $6.88 \times 10^{-1}$ | $3.26 \times 10^{-09}$ |
| *UBASH3B* | 11 | $2.79 \times 10^{-3}$ | $8.10 \times 10^{-4}$ | $5.22 \times 10^{-3}$ | $3.26 \times 10^{-08}$ |
| *HNF1A* | 12 | $4.65 \times 10^{-16}$ | $4.56 \times 10^{-16}$ | $NA$ | $1.96 \times 10^{-13}$ |
| *LIPC* | 15 | $1.56 \times 10^{-76}$ | $1.20 \times 10^{-71}$ | $2.40 \times 10^{-4}$ | $7.77 \times 10^{-15}$ |
| *CETP* | 16 | $2.67 \times 10^{-55}$ | $9.12 \times 10^{-50}$ | $1.20 \times 10^{-4}$ | $5.00 \times 10^{-10}$ |
| *HPR* | 16 | $2.49 \times 10^{-32}$ | $9.71 \times 10^{-33}$ | $6.95 \times 10^{-1}$ | $2.37 \times 10^{-12}$ |
| *LIPG* | 18 | $1.08 \times 10^{-14}$ | $1.07 \times 10^{-14}$ | $NA$ | $4.31 \times 10^{-07}$ |
| *APOE* | 19 | $1.00 \times 10^{-300}$ | $1.00 \times 10^{-300}$ | $5.14 \times 10^{-2}$ | $7.77 \times 10^{-15}$ |
| *HNF4A* | 20 | $9.74 \times 10^{-21}$ | $5.40 \times 10^{-21}$ | $3.48 \times 10^{-1}$ | $1.98 \times 10^{-3}$ |

(D) TG: gene-level association testing

| hgID | chr | EPIC | | | MAGMA |
|---|---|---|---|---|---|
| *GCKR* | 2 | $1.34 \times 10^{-289}$ | $6.55 \times 10^{-291}$ | $3.80 \times 10^{-2}$ | $5.00 \times 10^{-10}$ |
| *RSPO3* | 6 | $3.44 \times 10^{-09}$ | $1.92 \times 10^{-09}$ | $3.31 \times 10^{-1}$ | $9.82 \times 10^{-6}$ |
| *MLXIPL* | 7 | $1.35 \times 10^{-125}$ | $4.82 \times 10^{-126}$ | $1.60 \times 10^{-1}$ | $5.00 \times 10^{-10}$ |
| *LPL* | 8 | $1.00 \times 10^{-300}$ | $1.00 \times 10^{-300}$ | $1.33 \times 10^{-58}$ | $7.61 \times 10^{-15}$ |
| *TRIB1* | 8 | $1.16 \times 10^{-47}$ | $6.61 \times 10^{-48}$ | $4.10 \times 10^{-1}$ | $5.00 \times 10^{-10}$ |
| *AKR1C4* | 10 | $1.99 \times 10^{-08}$ | $1.99 \times 10^{-08}$ | $NA$ | $3.48 \times 10^{-08}$ |
| *LIPC* | 15 | $5.72 \times 10^{-29}$ | $9.92 \times 10^{-30}$ | $1.95 \times 10^{-05}$ | $8.47 \times 10^{-14}$ |
| *FRMD5* | 15 | $2.58 \times 10^{-06}$ | $2.56 \times 10^{-11}$ | $6.01 \times 10^{-4}$ | $8.18 \times 10^{-09}$ |
| *CETP* | 16 | $1.99 \times 10^{-34}$ | $2.53 \times 10^{-37}$ | $7.48 \times 10^{-10}$ | $4.56 \times 10^{-14}$ |
| *PLA2G6* | 22 | $8.68 \times 10^{-5}$ | $8.54 \times 10^{-05}$ | $NA$ | $1.12 \times 10^{-06}$ |

(E) SCZ: gene-level association testing

| hgID | chr | EPIC | | | MAGMA |
|---|---|---|---|---|---|
| | | Joint | Common | Rare | |
| *FXR1* | 3 | $1.27 \times 10^{-17}$ | $2.99 \times 10^{-17}$ | $4.73 \times 10^{-1}$ | $1.83 \times 10^{-09}$ |
| *CNTN4* | 3 | $1.18 \times 10^{-2}$ | $1.22 \times 10^{-2}$ | $NA$ | $9.59 \times 10^{-09}$ |
| *HCN1* | 5 | $1.38 \times 10^{-08}$ | $4.74 \times 10^{-4}$ | $1.53 \times 10^{-1}$ | $1.39 \times 10^{-06}$ |
| *SNAP91* | 6 | $1.07 \times 10^{-14}$ | $9.93 \times 10^{-15}$ | $4.97 \times 10^{-1}$ | $2.41 \times 10^{-10}$ |
| *GRM3* | 7 | $1.84 \times 10^{-11}$ | $1.88 \times 10^{-10}$ | $3.74 \times 10^{-2}$ | $1.94 \times 10^{-5}$ |
| *DRD2* | 11 | $2.56 \times 10^{-17}$ | $1.81 \times 10^{-19}$ | $7.57 \times 10^{-2}$ | $3.04 \times 10^{-09}$ |
| *NRGN* | 11 | $2.48 \times 10^{-12}$ | $9.05 \times 10^{-13}$ | $9.80 \times 10^{-1}$ | $5.23 \times 10^{-11}$ |
| *ATP2A2* | 12 | $5.39 \times 10^{-11}$ | $1.67 \times 10^{-06}$ | $1.71 \times 10^{-1}$ | $5.87 \times 10^{-3}$ |
| *BCL11B* | 14 | $2.00 \times 10^{-2}$ | $2.02 \times 10^{-2}$ | $3.38 \times 10^{-3}$ | $3.74 \times 10^{-09}$ |
| *CHRNA3* | 15 | $1.64 \times 10^{-15}$ | $2.34 \times 10^{-15}$ | $2.44 \times 10^{-1}$ | $8.86 \times 10^{-10}$ |
| *CHRNB4* | 15 | $4.36 \times 10^{-11}$ | $1.70 \times 10^{-11}$ | $8.94 \times 10^{-1}$ | $1.15 \times 10^{-06}$ |
| *CACNA1I* | 22 | $4.51 \times 10^{-17}$ | $2.85 \times 10^{-05}$ | $4.41 \times 10^{-1}$ | $3.24 \times 10^{-13}$ |

(F) BIP: gene-level association testing

| hgID | chr | EPIC | | | MAGMA |
|---|---|---|---|---|---|
| | | Joint | Common | Rare | |
| *PLEKHO1* | 1 | $7.54 \times 10^{-6}$ | $6.67 \times 10^{-07}$ | $3.93 \times 10^{-1}$ | $1.31 \times 10^{-06}$ |
| *TRANK1* | 3 | $1.39 \times 10^{-19}$ | $2.24 \times 10^{-17}$ | $1.88 \times 10^{-1}$ | $5.17 \times 10^{-08}$ |
| *ITIH1* | 3 | $4.07 \times 10^{-06}$ | $1.13 \times 10^{-4}$ | $3.73 \times 10^{-1}$ | $1.39 \times 10^{-06}$ |
| *FSTL5* | 4 | $5.81 \times 10^{-08}$ | $1.54 \times 10^{-07}$ | $7.47 \times 10^{-2}$ | $4.18 \times 10^{-2}$ |
| *RIMS1* | 6 | $2.62 \times 10^{-06}$ | $2.87 \times 10^{-06}$ | $2.35 \times 10^{-1}$ | $8.49 \times 10^{-4}$ |
| *TFAP2B* | 6 | $3.20 \times 10^{-06}$ | $5.91 \times 10^{-2}$ | $1.49 \times 10^{-1}$ | $1.28 \times 10^{-07}$ |
| *FADS2* | 11 | $2.66 \times 10^{-06}$ | $3.71 \times 10^{-06}$ | $2.47 \times 10^{-3}$ | $6.76 \times 10^{-08}$ |
| *NCAN* | 19 | $1.15 \times 10^{-08}$ | $4.82 \times 10^{-06}$ | $3.23 \times 10^{-1}$ | $3.66 \times 10^{-5}$ |

(G) SCZBIP: gene-level association testing

| hgID | chr | EPIC | | | MAGMA |
|---|---|---|---|---|---|
| | | Joint | Common | Rare | |
| *PLEKHO1* | 1 | $1.42 \times 10^{-17}$ | $1.32 \times 10^{-17}$ | $NA$ | $7.93 \times 10^{-12}$ |
| *TRANK1* | 3 | $1.05 \times 10^{-19}$ | $1.04 \times 10^{-19}$ | $2.30 \times 10^{-2}$ | $2.37 \times 10^{-13}$ |
| *ITIH1* | 3 | $2.37 \times 10^{-08}$ | $1.98 \times 10^{-08}$ | $1.21 \times 10^{-1}$ | $2.32 \times 10^{-09}$ |
| *FXR1* | 3 | $4.75 \times 10^{-08}$ | $3.01 \times 10^{-08}$ | $9.59 \times 10^{-1}$ | $4.83 \times 10^{-06}$ |
| *CNTN4* | 3 | $1.36 \times 10^{-2}$ | $1.87 \times 10^{-2}$ | $5.88 \times 10^{-2}$ | $2.69 \times 10^{-09}$ |
| *FSTL5* | 4 | $5.54 \times 10^{-07}$ | $8.86 \times 10^{-07}$ | $6.88 \times 10^{-2}$ | $1.61 \times 10^{-3}$ |
| *HCN1* | 5 | $1.37 \times 10^{-13}$ | $1.37 \times 10^{-13}$ | $2.53 \times 10^{-1}$ | $3.07 \times 10^{-07}$ |
| *SNAP91* | 6 | $2.82 \times 10^{-20}$ | $2.09 \times 10^{-20}$ | $2.91 \times 10^{-1}$ | $8.84 \times 10^{-12}$ |
| *TFAP2B* | 6 | $4.36 \times 10^{-06}$ | $4.34 \times 10^{-06}$ | $NA$ | $6.72 \times 10^{-08}$ |
| *GRM3* | 7 | $6.16 \times 10^{-11}$ | $4.92 \times 10^{-11}$ | $3.32 \times 10^{-1}$ | $4.63 \times 10^{-07}$ |
| *ANK3* | 10 | $1.66 \times 10^{-12}$ | $3.01 \times 10^{-12}$ | $5.23 \times 10^{-2}$ | $1.81 \times 10^{-5}$ |
| *DRD2* | 11 | $7.23 \times 10^{-17}$ | $7.07 \times 10^{-17}$ | $NA$ | $7.50 \times 10^{-07}$ |
| *NRGN* | 11 | $9.57 \times 10^{-10}$ | $9.44 \times 10^{-10}$ | $NA$ | $5.62 \times 10^{-08}$ |
| *BCL11B* | 14 | $5.32 \times 10^{-3}$ | $6.23 \times 10^{-3}$ | $2.10 \times 10^{-1}$ | $2.55 \times 10^{-12}$ |
| *CHRNA3* | 15 | $1.44 \times 10^{-15}$ | $6.02 \times 10^{-16}$ | $5.54 \times 10^{-1}$ | $3.90 \times 10^{-09}$ |
| *CHRNB4* | 15 | $1.51 \times 10^{-14}$ | $2.56 \times 10^{-14}$ | $6.69 \times 10^{-2}$ | $2.66 \times 10^{-07}$ |
| *GRIN2A* | 16 | $1.13 \times 10^{-06}$ | $2.56 \times 10^{-06}$ | $2.93 \times 10^{-2}$ | $5.12 \times 10^{-08}$ |
| *CACNA1I* | 22 | $5.94 \times 10^{-08}$ | $4.12 \times 10^{-08}$ | $7.65 \times 10^{-1}$ | $1.09 \times 10^{-12}$ |

(H) T2Db: gene-level association testing

| hgID | chr | EPIC | | | MAGMA |
|---|---|---|---|---|---|
| | | Joint | Common | Rare | |
| *GCKR* | 2 | $2.61 \times 10^{-25}$ | $3.66 \times 10^{-20}$ | $8.12 \times 10^{-2}$ | $1.29 \times 10^{-11}$ |
| *GRB14* | 2 | $1.29 \times 10^{-09}$ | $8.25 \times 10^{-10}$ | $4.51 \times 10^{-1}$ | $6.15 \times 10^{-4}$ |
| *PPARG* | 3 | $4.55 \times 10^{-68}$ | $5.68 \times 10^{-66}$ | $3.09 \times 10^{-05}$ | $2.33 \times 10^{-13}$ |
| *SCD5* | 4 | $9.21 \times 10^{-2}$ | $2.10 \times 10^{-1}$ | $4.84 \times 10^{-1}$ | $1.67 \times 10^{-06}$ |
| *PAM* | 5 | $1.46 \times 10^{-24}$ | $2.18 \times 10^{-24}$ | $6.87 \times 10^{-2}$ | $2.02 \times 10^{-13}$ |
| *SLC30A8* | 8 | $6.65 \times 10^{-65}$ | $9.53 \times 10^{-75}$ | $4.31 \times 10^{-1}$ | $2.15 \times 10^{-08}$ |
| *NEUROG3* | 10 | $3.79 \times 10^{-19}$ | $6.35 \times 10^{-18}$ | $2.98 \times 10^{-3}$ | $1.02 \times 10^{-09}$ |
| *KCNQ1* | 11 | $8.00 \times 10^{-114}$ | $3.23 \times 10^{-113}$ | $NA$ | $2.57 \times 10^{-24}$ |
| *INS* | 11 | $2.00 \times 10^{-21}$ | $5.77 \times 10^{-20}$ | $5.64 \times 10^{-2}$ | $5.02 \times 10^{-06}$ |
| *HNF1A* | 12 | $6.09 \times 10^{-30}$ | $1.22 \times 10^{-29}$ | $1.32 \times 10^{-3}$ | $1.31 \times 10^{-10}$ |
| *HNF4A* | 20 | $1.48 \times 10^{-54}$ | $4.10 \times 10^{-54}$ | $1.85 \times 10^{-1}$ | $4.60 \times 10^{-09}$ |

**Table S3. Top three tissue types identified using rare variants only in the GTEx bulk RNA-seq data.** We considered rare variants only and recovered the gene-level chi-square association test statistic from the burden test. Gene-gene correlations were retrieved from the joint analysis of common and rare variants with sliding windows. We identified top three tissue types using the framework of tissue-specific enrichment analysis. The asterisk (*) indicates statistical significance at the significance level 0.05 with Bonferroni correction.

| Trait | Top 1 relevant tissue for rare variants | Top 2 relevant tissue for rare variants | Top 3 relevant tissue for rare variants |
|---|---|---|---|
| LDL | Liver* | Small intestine | Colon transverse |
| HDL | Liver | Adrenal gland | Spleen |
| TC | Liver* | Small intestine | Colon transverse |
| TG | Liver | Thyroid | Pituitary |
| SCZ | Brain cerebellum | Brain cerebellar hemisphere | Heart left ventricle |
| BIP | Brain frontal cortex* | Brain cortex | Brain anterior |
| SCZBIP | Brain frontal cortex* | Brain cortex* | Brain anterior* |
| T2Db | Brain frontal cortex* | Brain cortex* | Brain cerebellar hemisphere* |

**Table S4: Number of common and rare variants from GWAS summary statistics with different thresholds.** We report the number of common and rare SNPs / the number of genes (average number of SNPs per gene) with different MAF and MAC thresholds. For rare variants, the upper bound of inclusion is controlled by MAF while the lower bound is determined by MAC. Rare variants with MAC less than 20 are removed from analysis by default.

**(A)** Number of rare variants with different thresholds of MAF (upper bound)

| Trait | MAF <= 1% | MAF <= 0.1% |
|---|---|---|
| LDL | 97,915 / 20,187 (5.5) | 90,528 / 19,550 (5.2) |
| HDL | 99,189 / 20,283 (5.6) | 90,918 / 19,575 (5.3) |
| TC | 98,930 / 20,315 (5.5) | 90,759 / 19,617 (5.2) |
| TG | 98,125 / 20,128 (5.5) | 90,548 / 19,476 (5.3) |
| SCZ | 1,959,467 / 30,935 (73.0) | 550,279 / 29,784 (21.3) |
| BIP | 1,487,888 / 30,878 (55.6) | 504 / 550 (1.1) |
| SCZBIP | 27,748 / 10,910 (2.9) | 0 / 0 (0) |
| T2Db | 3,224,387 / 30,991 (120.3) | 1,868,536 / 30,919 (69.6) |

**(B)** Number of rare variants with different thresholds of MAF (upper bound) and MAC filtering (lower bound)

| Trait | MAF <= 1% | MAF <= 0.1% |
|---|---|---|
| LDL | 44,412 / 3,164 (14.8) | 0 / 0 (0) |
| HDL | 47,651 / 3,437 (14.7) | 0 / 0 (0) |
| TC | 47,033 / 3,406 (14.6) | 0 / 0 (0) |
| TG | 44,812 / 3,204 (14.8) | 0 / 0 (0) |
| SCZ | 1,959,449 / 30,905 (73.1) | 548,393 / 28,364 (22.2) |
| BIP | 1,487,888 / 30,878 (55.6) | 503 / 549 (1.1) |
| SCZBIP | 27,748 / 10,910 (2.9) | 0 / 0 (0) |
| T2Db | 3,224,317 / 30,918 (120.5) | 1,867,997 / 30,613 (70.2) |

**(C)** Number of pruned-in common variants with different thresholds of MAF (lower bound)

| Trait | MAF > 1% | MAF > 0.1% |
|---|---|---|
| LDL | 221,998 / 27,288 (9.2) | 225,222 / 27,337 (9.3) |
| HDL | 223,456 / 27,320 (9.2) | 227,066 / 27,378 (9.3) |
| TC | 223,429 / 27,322 (9.2) | 226,991 / 27,379 (9.3) |
| TG | 222,154 / 27,290 (9.2) | 225,444 / 27,341 (9.3) |
| SCZ | 1,114,970 / 31,030 (41.1) | 1,991,247 / 31,086 (73.5) |
| BIP | 1,086,963 / 31,030 (40.1) | 1,962,422 / 31,096 (72.5) |
| SCZBIP | 1,087,425 / 30,980 (40.2) | 1,223,597 / 31,003 (45.2) |
| T2Db | 1,068,216 / 30,792 (39.7) | 2,052,221 / 30,935 (76.3) |

### Supplementary Figures

**Figure S1. Gene-level quantile-quantile plots for eight GWAS traits.** EPIC achieves higher power than MAGMA in the gene-level association test.

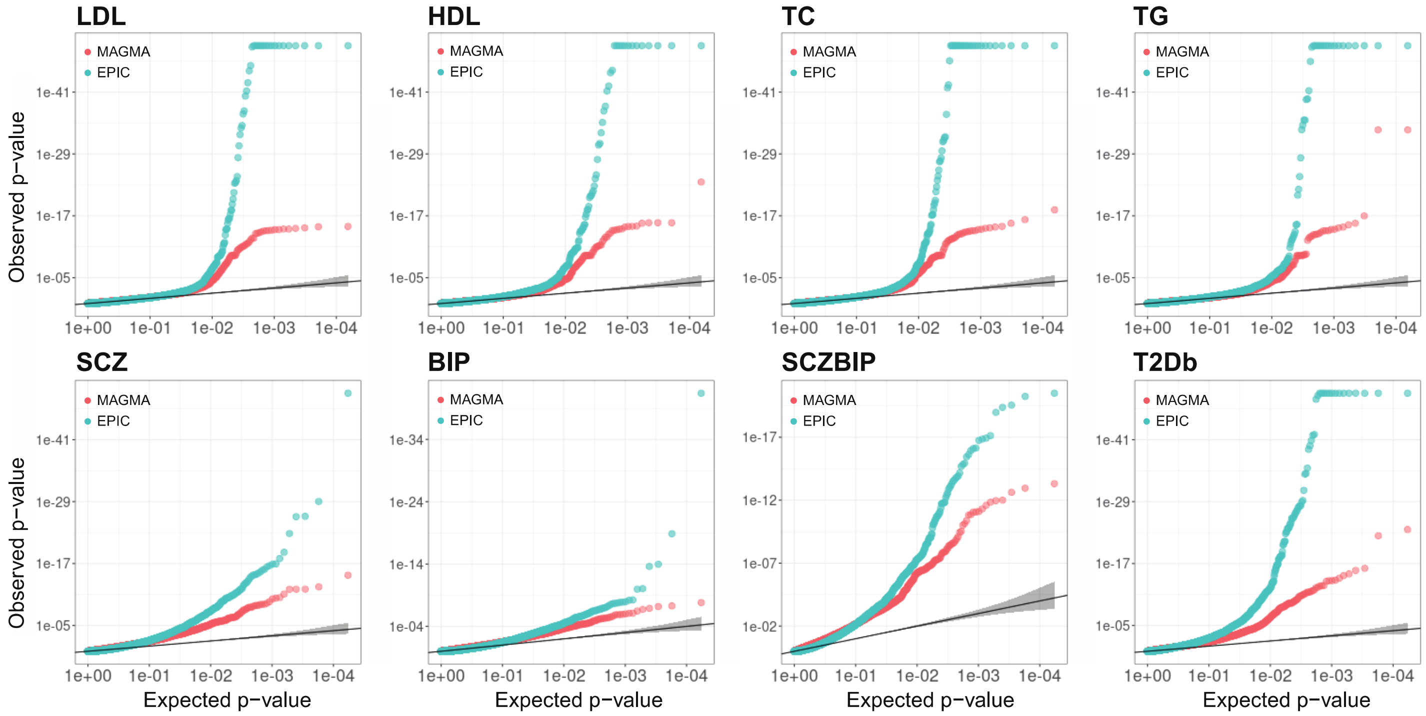

**Figure S2. Venn diagram of significant genes associated with eight GWAS traits by EPIC and MAGMA.** EPIC detected more significantly associated genes compared to MAGMA.

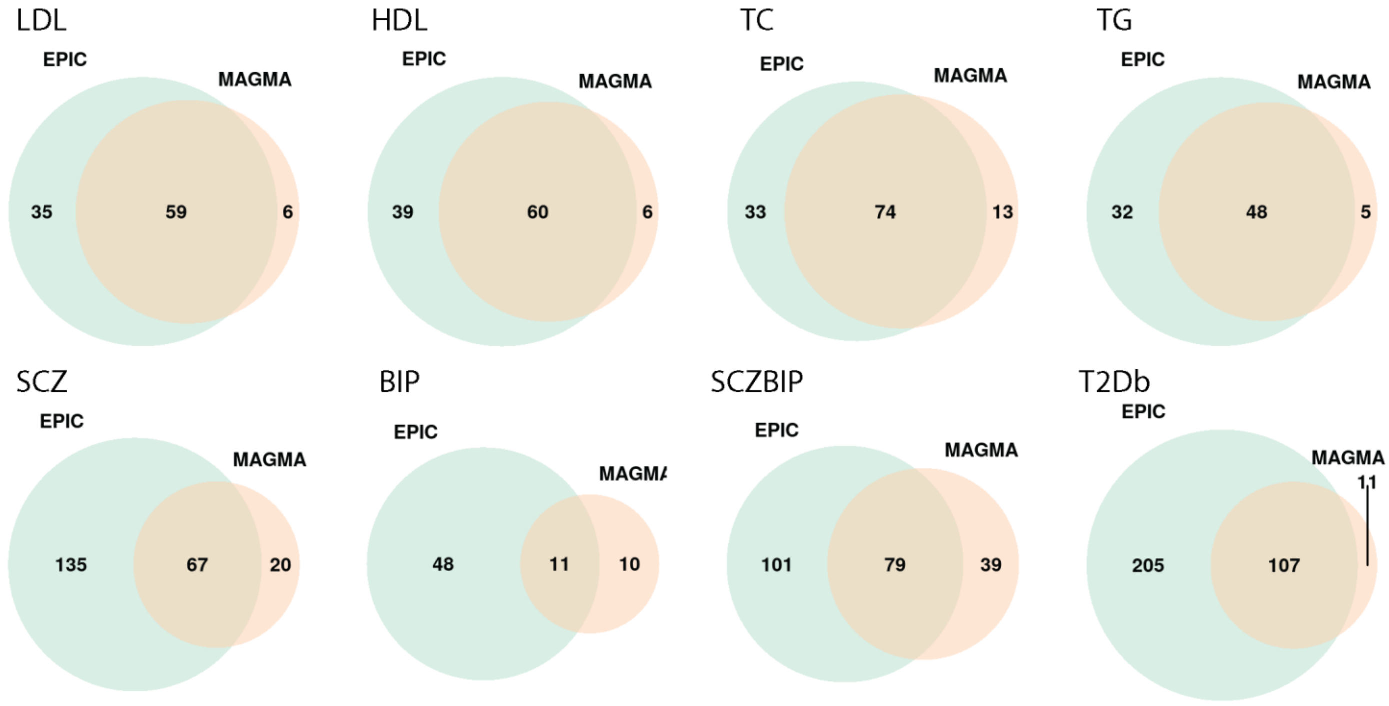

**Figure S3: Gene-level association test for housekeeping genes in the analysis of trait-relevant tissue identification using GTEx bulk RNA-seq data.** Boxplots of –log10(p-value). A final set of 8,708 genes are retained in the analysis, where there are 52 out of 2,088 overlapped housekeeping genes.

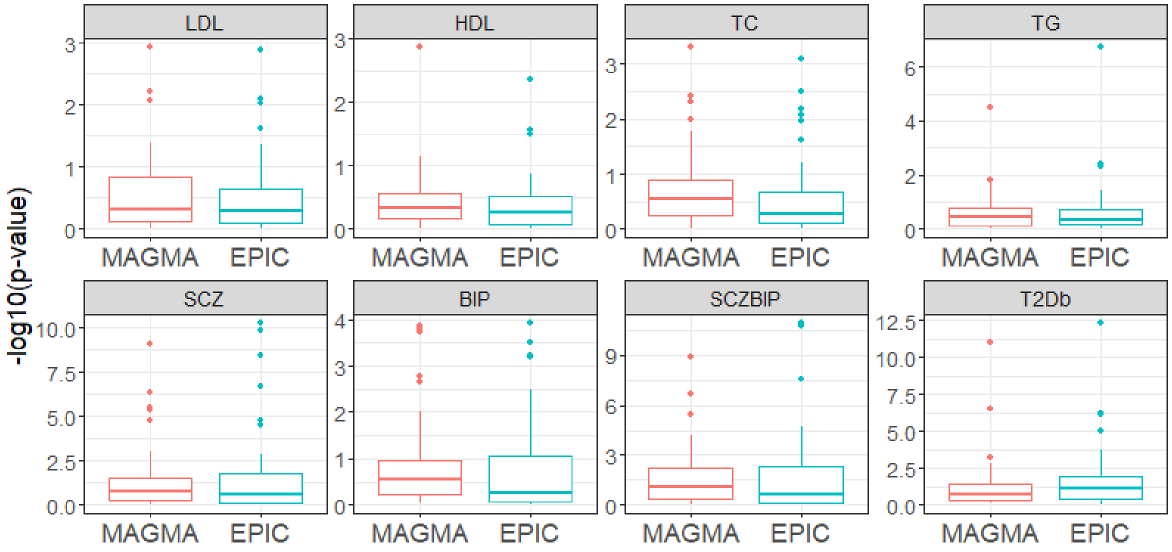

**Figure S4. UMAP plots of pancreatic islet scRNA-seq datasets.** (A) Baron et al. [8]: UMAP embedding of 8,569 InDrop single-cell profiles from three healthy donors; (B) Segerstolpe et al. [9]: UMAP embedding of 1,068 Smart-seq2 single-cell profiles from six healthy donors.

(A)

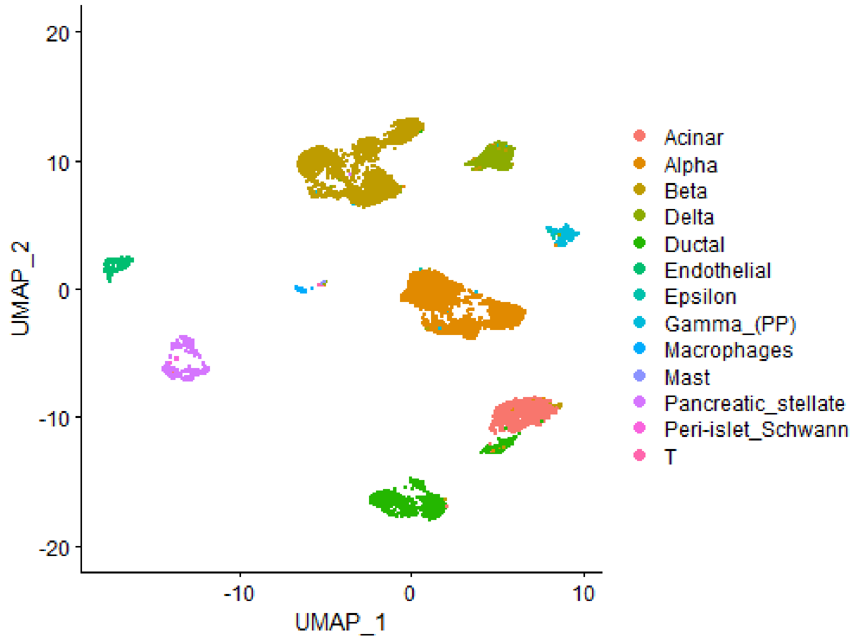

(B)

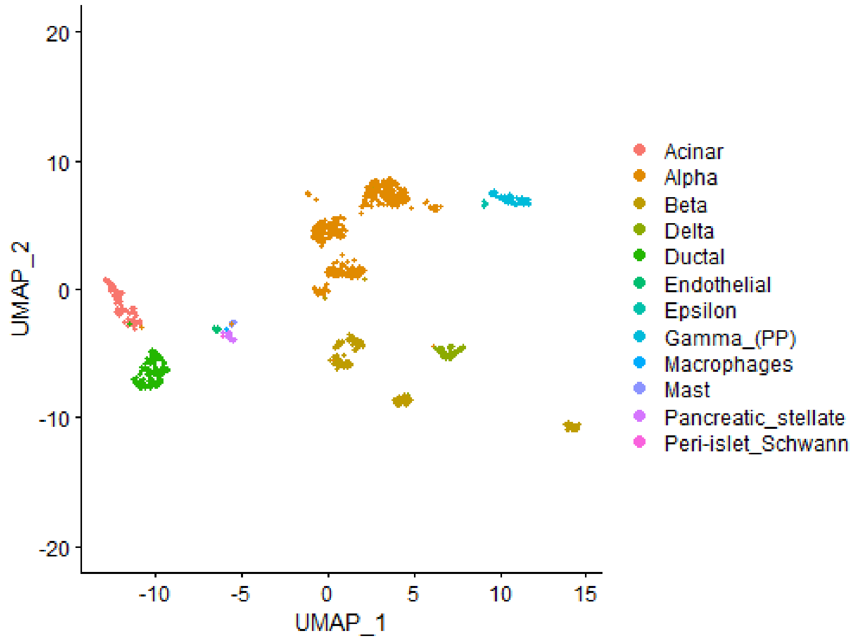

**Figure S5. Validation strategies of cell-type enrichment results for schizophrenia.** Volcano plot for 287 differentially expressed (DE) genes that were reported from an independent case versus control study for schizophrenia using bulk RNA-seq.

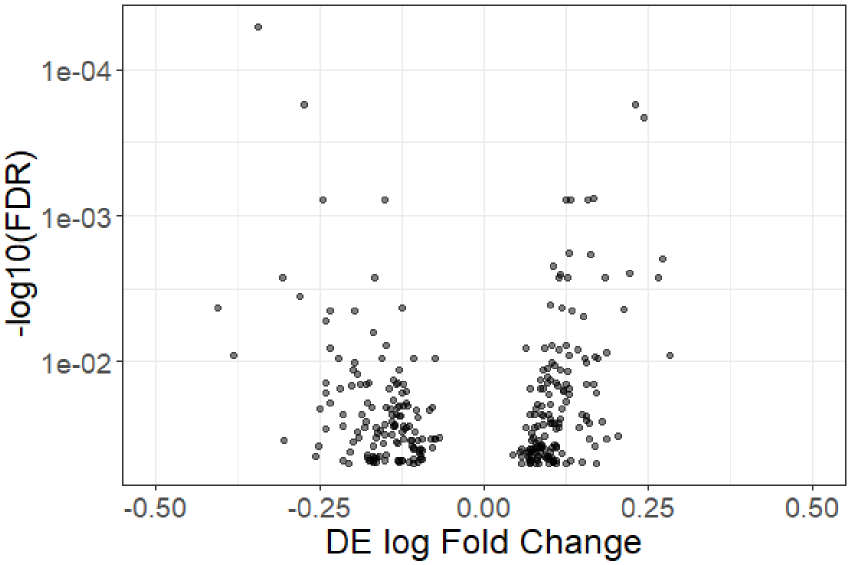

**Figure S6. Comparison of gene-based *p*-value with different sliding window sizes.** We evaluated the effects of different choices of sliding window size on the gene-level *p*-values. A final set of 8,708 genes are retained in the analysis of the GTEx bulk RNA-seq dataset. The results were not substantially altered by sliding window size while the computational burden was considerably increased. To account for gene-gene correlations and improve computational efficiency, the sliding window size is set to be 10.

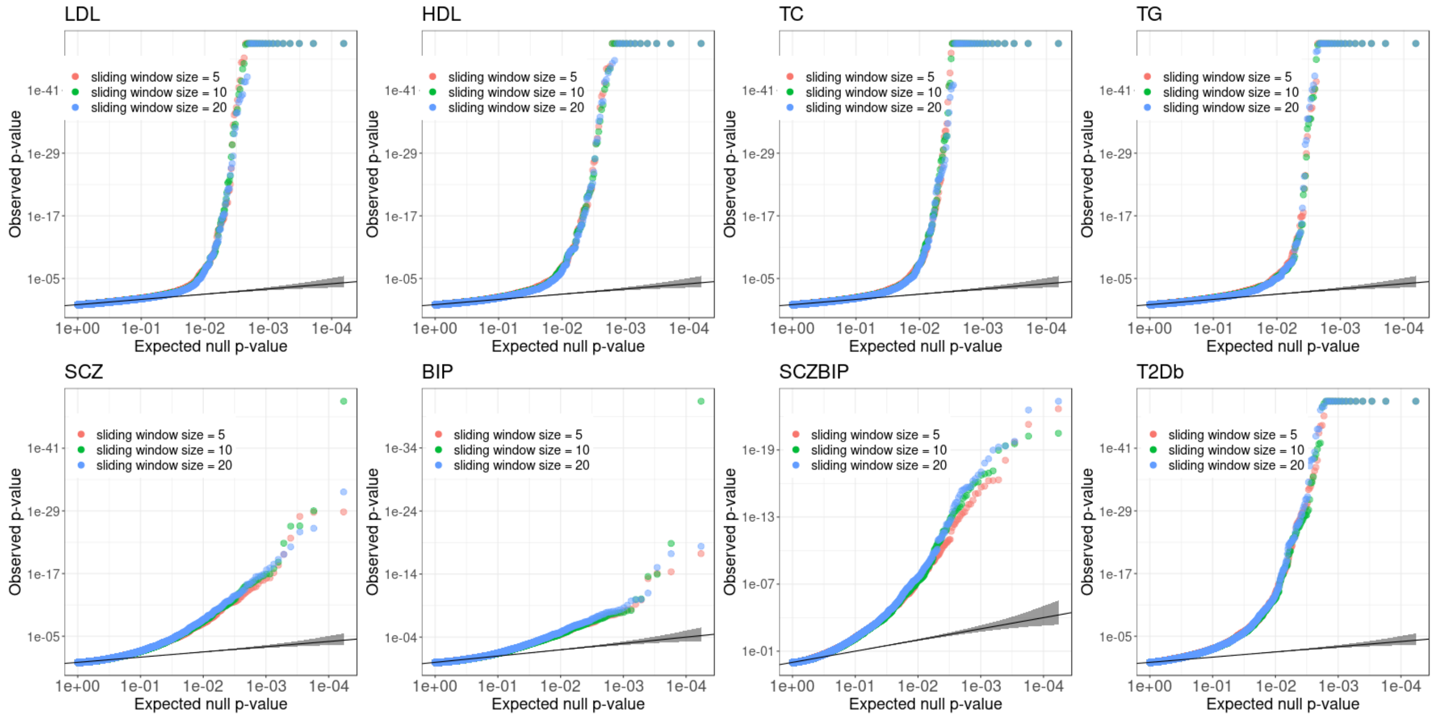

**Figure S7. Enrichment results comparison using different MAF cutoffs.** Tissue- or cell-type-specific enrichment results by EPIC (-log(p-value)) with MAF=1% and MAF=0.1% cutoffs. EPIC is robust to the choice of MAF thresholds.

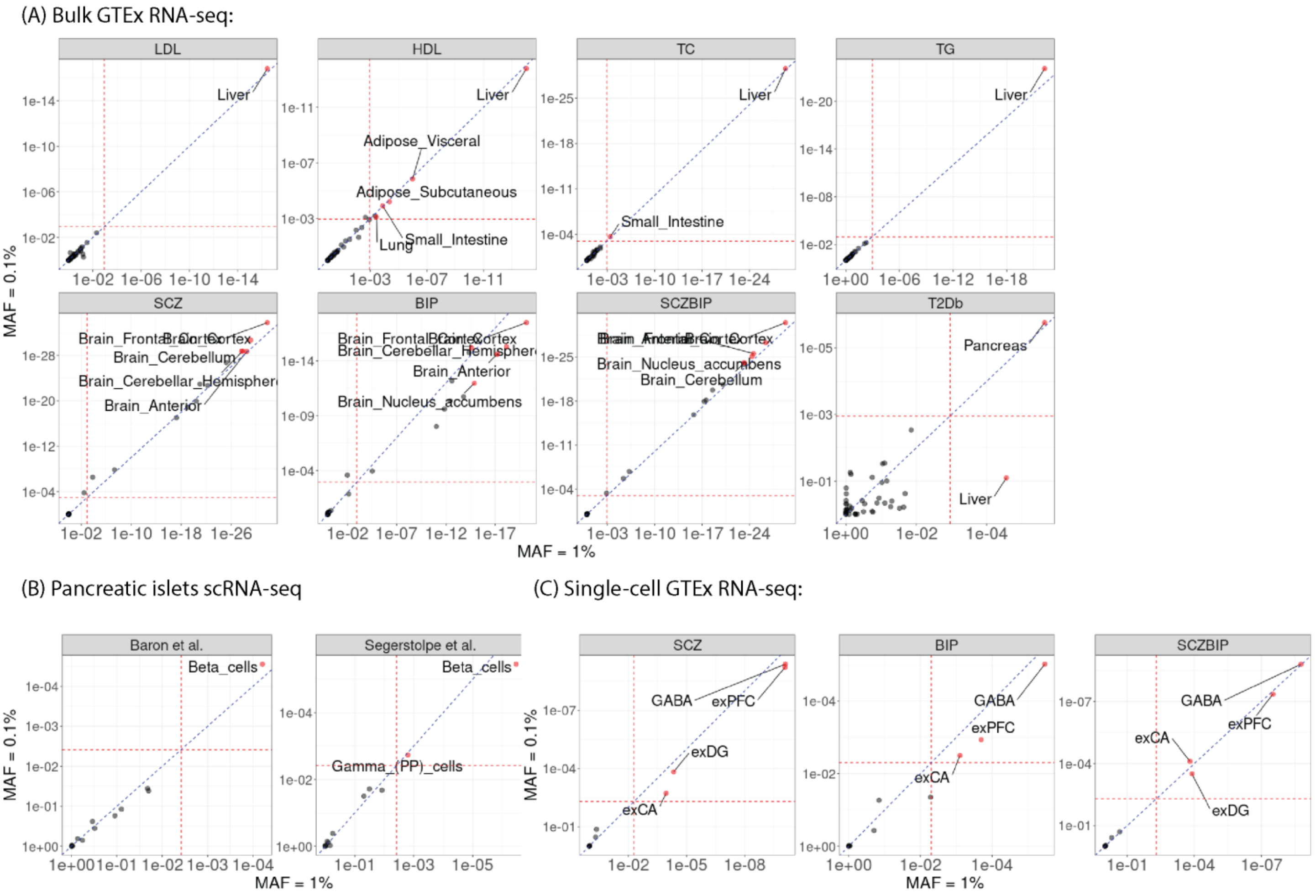

**Figure S8. Effect of LD pruning thresholds for common variants.** The left panel shows the percentage of gene-specific LD pruning thresholds on the genome-wide scale. The right panel shows the number of common SNPs per gene across different LD pruning thresholds for genes that need a second-round pruning. The threshold of 0.8 corresponds to the majority of genes with only one-round of pruning. The thresholds less than 0.8 correspond to the remaining genes that need a second round of pruning.

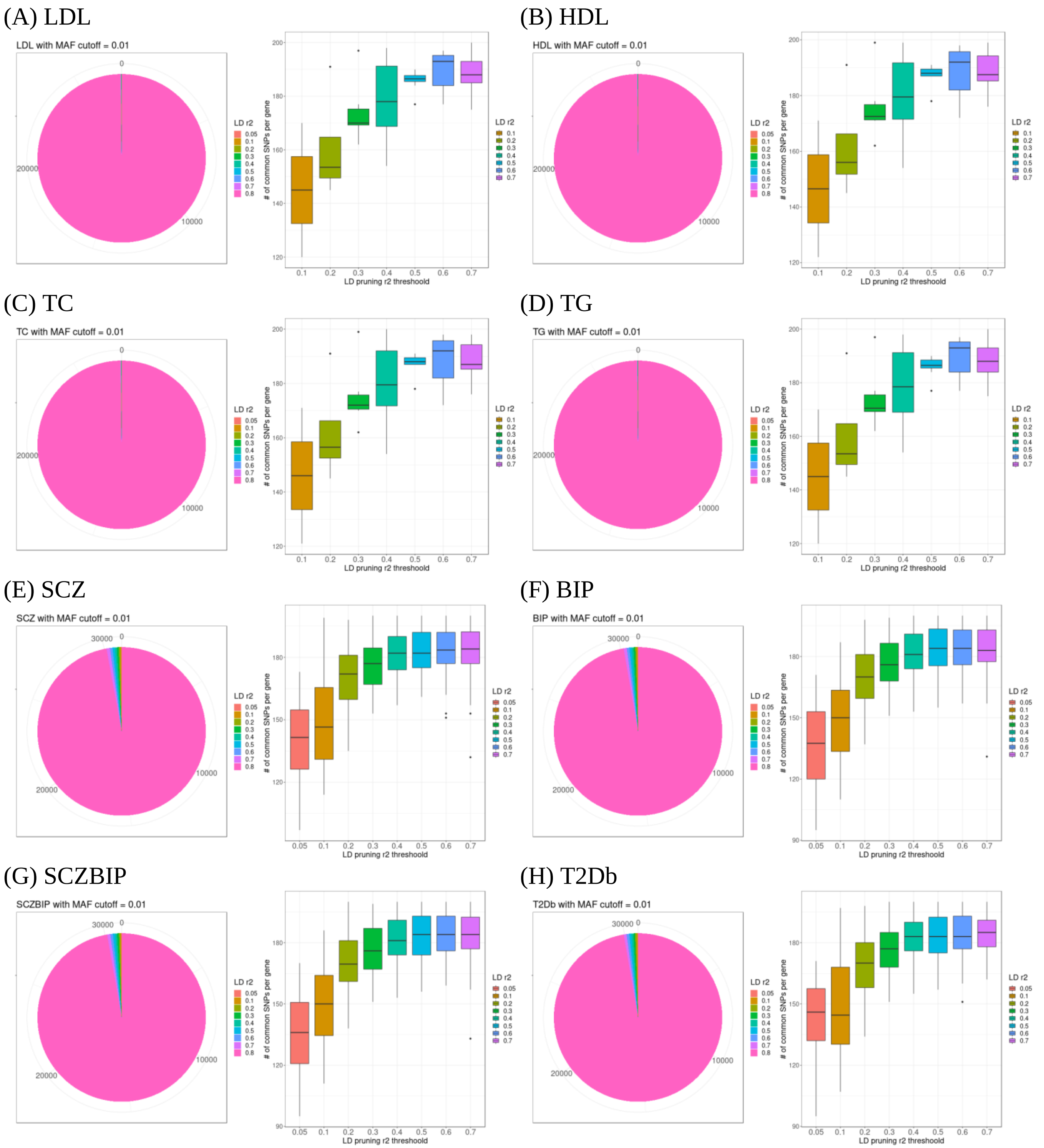

### References

1. Willer CJ*, et al.* Discovery and refinement of loci associated with lipid levels. *Nat Genet* **45**, 1274-1283 (2013).

2. Schizophrenia Working Group of the Psychiatric Genomics C. Biological insights from 108 schizophrenia-associated genetic loci. *Nature* **511**, 421-427 (2014).

3. Pardinas AF*, et al.* Common schizophrenia alleles are enriched in mutation-intolerant genes and in regions under strong background selection. *Nat Genet* **50**, 381-389 (2018).

4. Stahl EA*, et al.* Genome-wide association study identifies 30 loci associated with bipolar disorder. *Nat Genet* **51**, 793-803 (2019).

5. Bipolar D, Schizophrenia Working Group of the Psychiatric Genomics Consortium. Electronic address drve, Bipolar D, Schizophrenia Working Group of the Psychiatric Genomics C. Genomic Dissection of Bipolar Disorder and Schizophrenia, Including 28 Subphenotypes. *Cell* **173**, 1705-1715 e1716 (2018).

6. Mahajan A*, et al.* Fine-mapping type 2 diabetes loci to single-variant resolution using high-density imputation and islet-specific epigenome maps. *Nat Genet* **50**, 1505-1513 (2018).

7. Consortium GT. The GTEx Consortium atlas of genetic regulatory effects across human tissues. *Science* **369**, 1318-1330 (2020).

8. Baron M*, et al.* A Single-Cell Transcriptomic Map of the Human and Mouse Pancreas Reveals Inter- and Intra-cell Population Structure. *Cell Syst* **3**, 346-360 e344 (2016).

9. Segerstolpe A*, et al.* Single-Cell Transcriptome Profiling of Human Pancreatic Islets in Health and Type 2 Diabetes. *Cell Metab* **24**, 593-607 (2016).

10. Habib N*, et al.* Massively parallel single-nucleus RNA-seq with DroNc-seq. *Nat Methods* **14**, 955-958 (2017).
